# PolliCrop: A high-throughput computer vision pipeline for pollinator monitoring in agroecosystems

**DOI:** 10.64898/2026.07.08.737348

**Authors:** Stan Chabert, Jordan Bernigaud-Samatan, Benjamin K. Blackman, Nicolas Blanchet, Olivier Catrice, Cécile Donnadieu, Marianne Gani, Rémi Grousset, Salena Husband, Guillaume Tueux, Silvio Erler, Nicolas B. Langlade

**Affiliations:** Univ Toulouse, CNRS, INRAE, LIPME, Castanet-Tolosan, France; Unité Expérimentale d’Agroécologie et de Phénotypage des Cultures (UE APC), INRAE, Castanet-Tolosan, France; Department of Plant & Microbial Biology, University of California, Berkeley, USA; Department of Integrative Biology, University of California, Berkeley, USA; Institute for Bee Protection, Julius Kühn Institute (JKI) – Federal Research Centre for Cultivated Plants, Braunschweig, Germany; Zoological Institute, Technische Universität Braunschweig, Braunschweig, Germany

**Keywords:** Crops, Deep learning, Flower attractiveness, Flower-visiting insects, Monitoring, Phenotyping, Sunflower

## Abstract

Flower-visiting insect populations are declining since the 1990s, especially because of the decrease of floral resources in agricultural settings. Mass flowering crops can help increase resource availability, and plant breeding can be directed towards selecting varieties attracting more flower-visiting insects. This requires the implementation of an automated high-throughput phenotyping tool for assessing the attractiveness of plant genotypes to flower-visiting insects. In this study, (*i*) we present a procedure to take standardized images of sunflower heads with camera traps continuously at day and night in the field; (*ii*) we trained two versions of a deep learning model, named PolliCrop, to automatically detect and identify the three insect classes visiting the most sunflower (non-*Bombus* bees, bumble bees, lepidopterans); (*iii*) we assessed and validated the ability of PolliCrop to correctly predict the true visitation frequencies of the insect classes on three sunflower genotypes; (*iv*) we presented two statistical approaches to compare the insect visitation frequencies between plant genotypes, one including weather variables, and the other one without. One PolliCrop version yielded satisfying performance to correctly detect the three insect classes. In particular, it correctly predicted the insect visitation frequencies on two sunflower genotypes in a range of ±10%. The other PolliCrop version can be useful in certain contexts of images and objectives. PolliCrop can be extended in the future to other crop species by training PolliCrop on new images captured in these crops. The field experimental design to set up for comparing the attractiveness between genotypes is also discussed.

## 1. Introduction

Global changes in land utilization have converted natural habitats to cultivated areas and consequently been a key driver of declines in the diversity and abundance of flower-visiting insect communities. These declines have been particularly marked since the 1990s, especially for wild and managed bees (Koh et al., 2016; Potts et al., 2016; Zattara & Aizen, 2021). Agricultural intensification, with the expansion of monocultures and the increasing use of pesticides, fertilizers, tillage, grazing, and mowing, has exacerbated these declines (Ollerton et al., 2014; Goulson et al., 2015; Grab et al., 2019). The flower-visiting insect communities are strongly impacted by the depletion, standardization, and impoverishment of floral resources, phenomena that are particularly important for pollen during spring and summer, and during the late-season months of summer for both pollen and nectar (Requier et al., 2015, 2017; Di Pasquale et al., 2016; Dolezal et al., 2019; Harris et al. 2024). Mass flowering crops can help increase resource availability in agricultural settings (Westphal et al., 2003; Holzschuh et al., 2013; Goulson et al., 2015), a noteworthy example of which is sunflower (*Helianthus annuus*) crop, blooming during the critical summer period of lowest flower resource availability (Requier et al., 2015, 2017; Husband et al., 2025).

Plant breeding can be directed towards selecting varieties that produce nectar and pollen in higher quantities and of higher quality to help support populations of flower-visiting insects (Prasifka et al., 2018; Harris and Ratnieks, 2026). Achieving this objective requires the implementation of automated, simple, and standardized high-throughput phenotyping tools for assessing the attractiveness of plant genotypes to flower-visiting insects (Pérez-Alfocea et al., 2024; Borghi et al., 2025). Insect visitation rates to flowers are traditionally assessed either by walking along plant transects and counting all the insects visiting the plants of the transects (Vaissière et al., 2011; Chabert et al., 2026), or by standing around a fixed area during a certain period of time and by counting all the insects visiting a focal chosen group of flowers during that time period (Fijen and Kleijn, 2017; Garibaldi et al., 2020). These manual observations are time-consuming, can be subject to observer bias, can deter insect visitations to flowers, can underestimate visitation rates, and, most importantly, sample only a fraction of the daily activity of insects as these observations usually occur during some limited daily time period (Steen, 2017; Pegoraro et al., 2020; Serra-Marin et al., 2025). Because these observations are generally conducted only during daylight hours, they necessarily miss potential activity made by insects during the night (Knop et al., 2018; Macgregor and Scott-Brown, 2020).

Automatic and remote monitoring of wildlife with cameras has existed since the 1960’s (Cutler and Swann, 1999). Historically, two types of devices were used to monitor vertebrate activity: 1) ‘time-lapse’ photography, i.e., when images are programmed to be automatically and regularly taken by the camera at a predefined time interval, or 2) ‘animal-triggered’ photography, i.e., when the image capture is triggered by an infrared (IR) sensor that detects changes in heat energy caused by a moving animal (Cutler and Swann, 1999; Swann et al., 2004). Time-lapse is appropriate when the probability for a target animal to move in front of the camera is high. The higher the probability, the more spaced out the image capture can be. Conversely, animal-triggering is more appropriate when this probability is low. However, IR-triggered cameras are not appropriate for invertebrates such as insects, since poikilotherms do not produce their own heat and are thus hardly detectable by IR sensors (Gardiner et al., 2026). Alternatively, some authors developed a system by coupling a digital video recorder with a video motion detection (VMD) sensor in which the recordings of short time-lapse with high frame rates are triggered by the movements of animals recorded between successive images (Steen, 2017; Barlow and O’Neill, 2020). This system is well adapted for monitoring insects visiting flowers (e.g., Steen and Aase, 2011; Steen, 2012), but it is frequently triggered by wind and not easily replicable on a large scale for high-throughput monitoring due to the cumbersome and expensive VMD sensor (Pegoraro et al., 2020). Lastly, Darras et al. (2024), Gardiner et al. (2026) or Smith et al. (2026) reused this system by replacing the VMD sensor by a light on-device deep learning insect recognition system, triggering autonomously image capture immediately after automatic insect recognition. This last system is not sensitive to wind and more adapted to routine, large scale monitoring.

Video recording to monitor flower-visiting insects in a continuous way started in the early 2000s. Sessions usually last from a few hours to 24 or 48 hours (e.g., Lortie et al., 2012; Edwards et al., 2015; Droissart et al., 2021; Wonderlin, 2024; Blareau et al., 2025). Videos enable the monitoring of all the insect activity occurring on flowers during the session time, including the rare events, without the inconvenience of carrying cumbersome VMD sensors in the field. However, visually analyzing the whole footage can be very tedious and may require reducing the reviewing to a subsample only (e.g., Blareau et al., 2025), or the use of programs to automatically detect motion events in video streams (Weinstein, 2015, 2018). Time-lapse image series and videos can both be recorded during the day or the night. For nighttime time-lapse photography, the field of view is illuminated with a flash when the images are captured (e.g., Suetsugu and Hayamizu, 2014; Alison et al., 2022; Shibata and Kudo, 2025). For nighttime video recording, the field of view is continuously illuminated with an IR light that does not scare away nocturnal insects (e.g., Droissart et al., 2021; Wonderlin, 2024; Blareau et al., 2025; Serra-Marin et al., 2025).

Continuous monitoring with videos or with animal-triggered photography enables the measurement of insect visitation rates to flowers, i.e., the total absolute number of insect visitations a flower received per unit of time. This number can then be used for crop pollination management (Garibaldi et al., 2020). On the other hand, time-lapse photography enables the measurement of a relative index of insect visitation frequency to flowers, i.e., the total number of flower-visiting insects counted in a given set of images divided by the number of images. This index can only be used to make comparisons, for instance between locations, time points, plant species, plant genotypes, etc. Overall, time-lapse photography is used more often than videos in studies monitoring wild animals, and most of the animals monitored are mammals while invertebrates are still a very small minority (Pollet et al., 2025). Compared to human observations, automatic monitoring with time-lapse or videos allows for the study of the whole daily activity of insects (e.g., Szabo, 1980; Stelzer and Chittka, 2010; Steen, 2017; Høye et al., 2021).

Both time-lapse and videos require a lot of human time and labor of reviewing after image capture for identifying insect species, the frequency of their presence in the images, tracking their trajectory, etc. Recent advances in artificial intelligence have led to the development of new deep learning models that automatically detect and identify insects on images up to a certain taxonomic rank or morphogroup (Høye et al., 2021; Chiranjeevi et al., 2025; Spiesman, 2026). Several authors have trained deep learning models to detect and identify flower-visiting insects, either in a single class (Bjerge et al., 2023b; Alex et al., 2025; Serra-Marin et al., 2025; Gardiner et al., 2026), at the insect order level (Stark et al., 2023; Ştefan et al., 2025), or at the finest possible morphogroup level (Spiesman et al., 2021; Bjerge et al., 2023a,c, 2024; Sittinger et al., 2024; Bernauer et al., 2026; Smith et al., 2026).

The aim of our study was to train a deep learning model, named PolliCrop, to automatically detect and identify the insects making daytime and nighttime visitations to sunflower heads from images captured with time-lapse camera traps installed in the field. The advantage of sunflowers bearing a single head is that we can monitor all the open flowers of the same plant with a single camera placed in front of the head. The model was trained to detect three morphogroups of the main insects visiting sunflower head: “non-*Bombus*” bees (including honey bees, large halictids or mining bees, and leafcutter bees), bumble bees and lepidopterans (including moths and butterflies). For an illustration, we tested the model on an image set generated to compare the differences in insect visitation frequency between three sunflower genotypes. Insects were also counted manually per class to assess how well the model correctly predicts the three insect morphogroups and the true insect visitation frequency index for each genotype. Two statistical approaches are presented, one including the weather variables, and the other one without, to test whether the weather effect on insect activity need to be corrected when comparing genotypes that do not bloom at the same time. The daily activity of each morphogroup is also described for each sunflower genotype to gain more ecological insights on temporal flower-insect interactions.

## 2. Materials and methods

### 2.1. Image collection

Images were collected with Wingscapes cameras (model WCT-00126, TimelapseCam Pro, Wingscapes, USA) mounted on tripods with the lens oriented towards the sunflower head (Fig. 1). Cameras were placed approximately 30-50 cm (depending on the head size) in front of the heads, one day before the heads were opened until the complete end of bloom, i.e., once the last florets located in the very center of the head were at least on their second day of pistillate stage. Images were automatically captured every 5 minutes during the whole blooming period of heads, including days and nights. The lens focus was adjusted at the beginning and then once a day until the end of image recording. The images captured by this camera model are normally illuminated with a LED flash, but the LEDs were covered with uncolored light-tight tape to avoid overexposing images captured at night and to avoid interference with night insect activity (see Fig. S1). Heads bloomed for ca. 6-12 days depending on the genotype and temperature, and the total number of images recorded could amount to ca. 3,000 images per head (i.e., 12 images per hour amounting to 288 images per day). The images had a resolution of 3008 × 1692 pixels (∼ 5 MP) and were stored in JPG format on SD cards of 32 GB, capable of storing over 30,000 images. For detailed information related to all the settings of the camera, see Supplementary data. Cameras are powered with batteries, making them suitable for autonomous monitoring *in situ* for approximately three weeks with our settings.

**Fig. 1.**
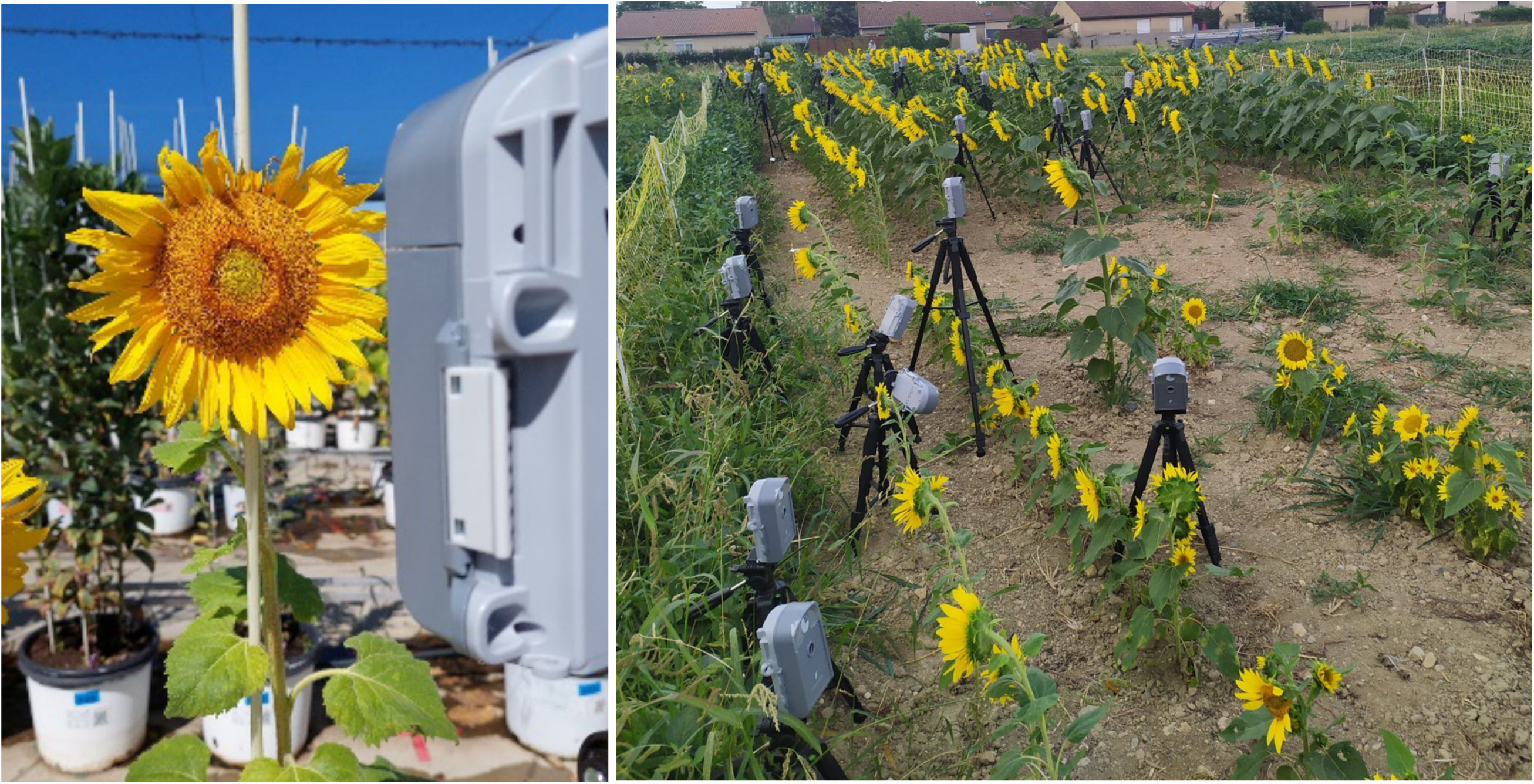
Left: Wingscapes camera positioned with the lens facing a sunflower head, on the Heliaphen platform. Right: Wingscapes cameras positioned on tripods in the field nursery of the INRAE site of Auzeville-Tolosane, France.

The images were collected in the INRAE research station of Auzeville-Tolosane, France (Latitude, 43.528; Longitude, 1.503), during the months of July, August and September of 2022, 2023 and 2024, both in the sunflower field nursery and on the Heliaphen platform (Fig. 1; Gosseau et al., 2019). Approximately 150,000 images were recorded on a variety of sunflower genotypes with diverse head morphologies, thus varying the image backgrounds for insect detection and identification.

### 2.2. Benchmark image set preparation

The 150,000 images were individually reviewed using Darwin V7 software package (Darwin V7 labs, London, UK). These images were initially sorted by (*i*) the presence of at least one insect on the image and (*ii*) the absence of any insects. Insect presence was considered regardless of where the insects were positioned in the images: on the head, leaves or stem, on the other sunflower plants in the background, or while flying. It was quite common to have two or more insects per image. But overall, there was an average of ca. 20% of images with insect presence. To define the insect classes, we focused in this study only on insect “pollinators” in a broad sense, i.e., insects that are known to actively forage for nectar and/or pollen in flowers and that can be broadly involved in pollination, depending on flower species. These taxa include Hymenoptera (including Vespidae and Anthophila, the clade name for “bees”; Branstetter et al., 2017; Murray et al., 2018), Lepidoptera (including butterflies and moths; Macgregor and Scott-Brown, 2020) and Diptera (Willmer et al., 2017). As these insect taxa are not necessarily pollinators of the flowers they visit, we refer to them with the broader terminology “flower-visiting insects” hereafter and in the whole manuscript (King et al., 2013). Fourteen morphogroups of flower-visiting insects were identified, plus three additional “morphogroups” of unidentified insects or other minority taxa (Table 1; Fig. 2). Only four morphogroups reached the minimum number of images (∼ 500) required to sufficiently train a deep learning model so that the model has the ability to detect and identify the morphogroup with enough performance: honey bees (*Apis mellifera*), large striped bees including Halictidae (halictids) and Andrenidae (mining bees), white end bumble bees (*Bombus* spp.) and moths (Heterocera). After training (see below), the deep learning model was able to detect and identify three classes of flower-visiting insects (Table 1): non-*Bombus* bees (including honey bees, large striped halictids and mining bees, and leafcutter bees); bumble bees (including white end bumble bees, red end bumble bees, brown bumble bees, and unidentified bumble bees); and lepidopterans (including moths and butterflies). Honey bees, large striped bees (halictids and mining bees), and leafcutter bees (*Megachile* spp.) were identified in the same class because of their similar size and appearance, with strips on the abdomen for all three morphogroups. All bumble bee morphogroups were identified in the same class because of their similar size, shape, and image texture. Lastly, butterflies and moths were identified in the same class because of their similar size, shape, and appearance of their wings, although moths are much more frequent than butterflies in the image set. Therefore, even if the model cannot distinguish between moths and butterflies for predictions, predictions are expected to mostly relate to moths, especially during the night.

**Fig. 2.**
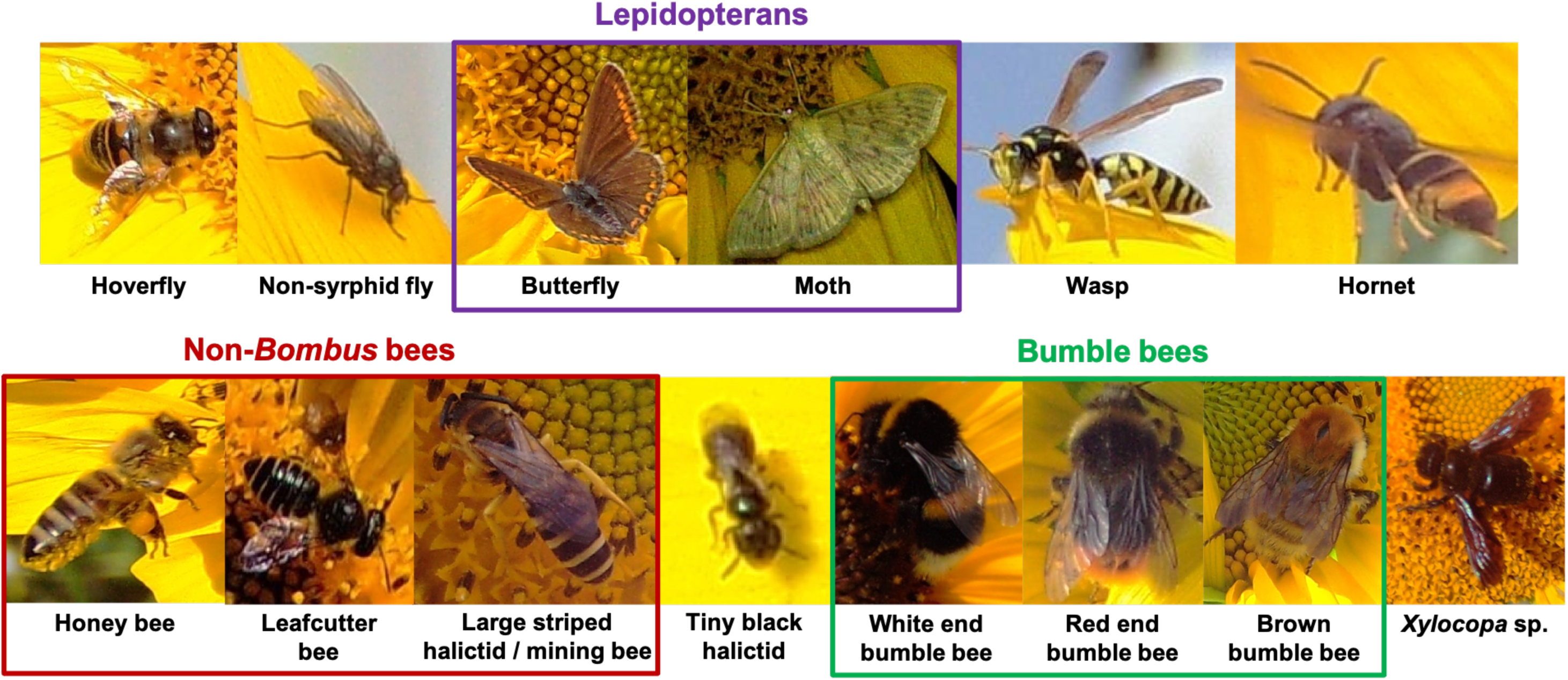
Morphogroups of flower-visiting insects identified on the 150,000 images reviewed of sunflower heads. The three colored boxes correspond to the three insect classes that the deep learning model was able to finally detect and identify after training: non-*Bombus* bees, bumble bees, and lepidopterans.

**Table 1.** Morphogroups of flower-visiting insects identified in the 150,000 images of sunflower heads reviewed, with the number of images in which each morphogroup was present.

| Insect order | Taxonomic subgroup | Other taxonomic subgroup | Morphogroup of flower-visiting insect | Number of images | Most probable insect species<sup>a</sup> |
| --- | --- | --- | --- | --- | --- |
| Diptera | Syrphidae |  | Hoverflies | 25 |  |
|  |  |  | Non-syrphid flies | 324 |  |
| Lepidoptera | Rhopalocera |  | Butterflies | 11 |  |
|  | Heterocera |  | **Moths** | **2570** |  |
| Hymenoptera | Vespidae |  | Wasps | 27 |  |
|  |  | *Vespa* spp. | Hornets | 9 |  |
|  | Anthophila (bees) | *Apis* | **Honey bees (*Apis mellifera*)** | **2942** | *Apis mellifera* |
|  |  | Megachilidae | Leafcutter bees (*Megachile* spp.) | 83 |  |
|  |  | Halictidae / Andrenidae | **Large striped halictids / mining bees** | **1168** | *Andrena flavipes*, *Halictus scabiosae* |
|  |  |  | Tiny black halictids | 207 | *Lasioglossum malachurum*, *L. subhirtum*, *L. pauxillum*, *Halictus tumulorum* |
|  |  | Bumble bees (*Bombus* spp.) | **White end *Bombus*** | **808** | *Bombus terrestris* agg., *B. hortorum*, *B. ruderatus* |
|  |  |  | Red end *Bombus* | 95 | *Bombus lapidarius* |
|  |  |  | Brown *Bombus* | 26 | *Bombus pascuorum* |
|  |  |  | Unidentified *Bombus* | 78 |  |
|  |  |  | *Xylocopa* spp. | 12 |  |
|  |  |  | Other Anthophila | 52 |  |
|  |  |  | Unidentified Anthophila | 116 |  |

In bold: morphogroups that reached the minimum number of images (ca. 500) required to sufficiently train the deep learning model so that the model has the ability to detect and identify the morphogroup with enough performance. The three colored boxes correspond to the three insect classes that the deep learning model was able to finally detect and identify after training: non-*Bombus* bees (in red), bumble bees (in purple), and lepidopterans (in green).
<sup>a</sup>Most probable bee species corresponding to the associated morphogroups, based on the species occurring at least 5 times in the databases of Rollin et al. (2015) and Aubouin et al. (2026) of bees foraging on sunflower in France.

Using Darwin V7 to prepare the benchmark image set to train the deep learning model, we assigned each flower-visiting insect present in a subset of the 150,000 images reviewed to one of the 17 morphogroups presented in Table 1. All of these insects were then cut out as polygons using Darwin V7’s annotation tool. The segmentation was adjusted with the brush tool to exclude the insect’s wings or legs to focus on the compact mass of the body, including the head, thorax and abdomen. The body is a constant component in the images, while wings and legs are not necessarily visible, or well visible, on all the images. However, the wings of lepidopterans were included in the segmentations since their wings are opaque, constitute an important part of lepidopterans’ body, and are therefore important for species identification. These segmentations were then converted to bounding box annotations for model training.

### 2.3. Training of the deep learning models PolliCrop with YOLO

The PolliCrop model was trained using the YOLO11x architecture (Redmon et al., 2016) through our public library *deepvisiontools* wrapper (version 0.2.0; https://forge.inrae.fr/ue-apc/librairies/python/deepvisiontools) built on Ultralytics (https://github.com/ultralytics/ultralytics). Two versions of PolliCrop were trained: yolov11_polinator3cls_04-2025 (hereafter named “PolliCrop1”), trained with 9,179 images, including 7,983 images presenting at least one insect belonging to one of the three classes identified previously (non-*Bombus* bees, bumble bees, lepidopterans; Table 1) and 1,196 (20%) “empty” images added to the training image set; and yolov11_polinator3cls_09-2025 (hereafter named “PolliCrop2”), trained with 14,448 images and including the same 7,983 images presenting insects as for PolliCrop1, but with the addition of 6,465 (45%) “empty” images distributed among the training, validation and testing image sets (see Table 2). The “empty” images correspond to images of sunflower heads, but without any insect on them. “Empty” images were added to decrease the number of false positive predicted. As insects were present only in a minority of images (∼ 20%), this low frequency can increase the risk for the model to “hallucinate” many insects in an image set (= ‘precision’ score, see below). An option to balance this risk is to include “empty” images, i.e., without any insect, in the benchmark image set. On the other hand, the inconvenience is to increase the risk for the model to miss many insects in an image set (= ‘recall’ score, see below).

**Table 2.** Assignment of the benchmark images to training, validation or testing.

| Image assignment | PolliCrop1 |  | PolliCrop2 |  | Both PolliCrop versions |  |  |  |
| --- | --- | --- | --- | --- | --- | --- | --- | --- |
|  | Total number of images | Number of empty<sup>a</sup> images | Total number of images | Number of empty<sup>a</sup> images | Number of images with insects | Number of non-*Bombus* bees | Number of bumble bees | Number of lepidopterans |
| Training | 6,124 (67%) | 1,196 (100%) | 8,547 (59%) | 3,619 (56%) | 4,928 (62%) | 2,982 (60%) | 675 (66%) | 1,613 (60%) |
| Validation | 1,715 (19%) | 0 | 2,921 (20%) | 1,206 (19%) | 1,715 (21%) | 1,066 (22%) | 201 (20%) | 650 (24%) |
| Testing | 1,340 (14%) | 0 | 2,980 (21%) | 1,640 (25%) | 1,340 (17%) | 886 (18%) | 148 (14%) | 426 (16%) |
| *Total* | *9,179* | *1,196* | *14,448* | *6,465* | *7,983* | *4,934* | *1,024* | *2,689* |

The percentages in brackets refer to the proportions compared to the total (last row).
<sup>a</sup>Images of sunflower heads without any insect.

The images were split into three groups: around 60% for the training image set, 20% for the validation image set, and 20% for the testing image set (Table 2). As multiple images were captured per plant, to avoid validating and testing the models on the same plants they were trained, all the images for a given plant were assigned to only one of the three groups of image sets (training, validation or test). Data preparation and modeling of the hyperparameters are detailed in the Supplementary data.

### 2.4. Calculations of performance scores to evaluate the models

After training, the PolliCrop models can predict the presence and location of an insect belonging to one of the three classes previously identified in images of sunflower heads. Non-*Bombus* bees are predicted in red boxes, bumble bees are predicted in purple boxes, and lepidopterans are predicted in green boxes (Fig. S2). Images presented in Fig. S2 correspond to true positives (TP), i.e., images where flower-visiting insects were correctly predicted by the models. True negatives (TN) are images for which the models correctly predicted the absence of any flower-visiting insect belonging to one of the classes previously presented. False positives (FP) are images for which one (or more) insect was “hallucinated” by the models (Fig. S3). False negatives (FN) are images for which the models failed to predict the presence of one (or more) insect (Fig. S4).

Performance scores can be calculated to evaluate the models on specific image sets, with the following formulas:

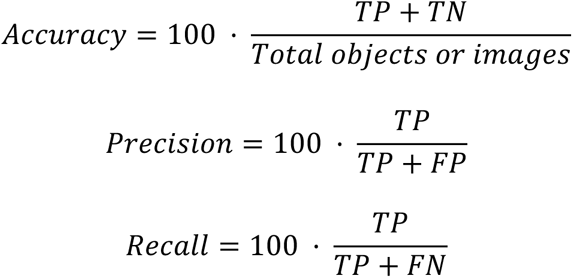

The ‘accuracy’ measures the percentage of images correctly predicted by the models. The ‘precision’ measures the percentage of images without “hallucinations” of insects by the models. The ‘recall’ measures the percentage of images for which the models did not miss insects. In this study, we calculated the scores at the object level for the evaluation from the testing image sets, or at the image level for other image sets, i.e., if the model did not predict the exact number of insects on a given image for a given insect class, the prediction was considered FP or FN for the image.

The two PolliCrop models were evaluated on two image sets. One image set, “22HP14”, was used to train the models. The images were captured in September 2022, on 14 sunflower plants on the Heliaphen platform (Gosseau et al., 2019). All the 8,970 images of this set were used to calculate the performance scores. The other image set, “24EV16”, correspond to images captured in August 2024 on plants of the Sunflower Extended Association Mapping population (Tueux et al., 2026). This population is composed of 435 cultivated inbred lines maximizing the sunflower genetic diversity of INRAE and USDA genetic resources (Mandel et al., 2011; Coque et al., 2008) and thus displaying a wide diversity of sunflower phenotypes. A sample of 1,024 images of this image set was used to calculate the performance scores. These images were not used to train the models.

To calculate the performance scores at the image level, these two image sets were first reviewed visually by humans, to count the number of insects present in the images belonging to each insect class (non-*Bombus* bees, bumble bees, lepidopterans). The two PolliCrop versions were then run on the two image sets (https://forge.inrae.fr/astr/public/pollicrop; for the procedure, see Supplementary data). The performance scores were calculated using the spreadsheet “Calculation_performance_scores” available at https://forge.inrae.fr/astr/public/pollicrop_toolbox.

### 2.5. Assessment of the ability of the two PolliCrop versions to correctly predict the insect visitation frequency of three sunflower genotypes

To assess the ability of the PolliCrop models to predict correctly the insect visitation frequency of sunflower heads, we used the image set ‘22HP14’ shortly described above. The aim of this experiment was to compare the insect visitation frequency of three sunflower research inbred lines, XRQ (n = 6), IR (n = 6) and CI (n = 3; Fig. S5). XRQ is the first sunflower genotype with a complete reference genome assembly and annotation (Badouin et al., 2017). The 15-L pots were filled with compost and regularly watered to avoid any water deficit until the end of the experiment. The plants bloomed from 5 to 16 September, 2022, and Wingscapes cameras were placed in front of the heads when they were blooming, capturing images every 5 minutes. A total of 8,970 images were recorded on which both versions of PolliCrop were run (https://forge.inrae.fr/astr/public/pollicrop; for the procedure, see Supplementary data), and all the images were also reviewed by a human to count the number of insects present belonging to each insect class (non-*Bombus* bees, bumble bees, lepidopterans). The PolliCrop models yielded a spreadsheet of data, named “PolliData”, giving the number of insects per class for each image by associating the image name. These data were first aggregated by hour to get the total sum of insect visitations per hour for each plant. This new dataset was named “PolliData_Hour”. Then, we connected these data to the genotype information, which was provided in a ‘MetaData’ file and merged it with the “PolliData_Hour” file using the software R (version 4.2.3; R Core team, 2023; R code available at https://forge.inrae.fr/astr/public/pollicrop_toolbox; see Supplementary data for procedure).

Before comparing the insect visitation frequencies assessed by the two versions of PolliCrop and the human eye for each sunflower genotype, we first investigated the day and night insect activity assessed from the counts made with the eye. Bees and bumble bees are expected to be active during the daylight hours (Szabo, 1980; Stelzer and Chittka, 2010; Steen, 2017; Høye et al., 2021), while moths are expected to be active during the night (Knop et al, 2018; Macgregor and Scott-Brown, 2020). This information may help to decrease the number of ‘zeros’, and thus data overdispersion for future analysis, by restricting the analysis to the hours of activity of each insect class. Early September in Auzeville-Tolosane, the sun rises around 07:30 (05:30 UTC) and sets around 20:15 (18:15 UTC). In parallel, the performance scores ‘precision’ and ‘recall’ of each PolliCrop version were calculated for each genotype and insect class.

### 2.6. Comparison of statistical models to assess the differences of insect visitation frequency between sunflower genotypes

Sunflower genotypes may bloom at different times because they may genetically differ in the thermal time regimes required to reach the blooming stage (Casadebaig et al., 2016). Consequently, comparisons of insect visitation frequency between genotypes may be made over different time intervals with different weather conditions. Yet insect activity is highly sensitive to the weather conditions. Bee foraging increases with increasing radiation and temperature (e.g., Szabo, 1980; Burrill and Dietz, 1981; Corbet et al., 1993; Vicens and Bosch, 2000; Clarke and Robert, 2018) until a thermal optimum beyond which it decreases (Colinet et al., 2015; Nielsen et al., 2017; Kenna et al., 2021; Jaboor et al., 2022), it decreases with increasing wind speed with a sigmoid shape (e.g., Pinzauti, 1986; Vicens and Bosch, 2000; Sanderson et al., 2015; Ngo et al., 2021), and it decreases with increasing rain intensity (Sanderson et al., 2015; Ngo et al., 2021). Lepidoptera foraging responds positively with increasing temperature as well (Knop et al, 2018). Therefore, it can be necessary to include the weather conditions in the statistical models to correctly compare the genotypes for insect visitation frequency. Two statistical models were compared for each insect class: one including the weather variables and one without.

To compare the statistical approaches, we used the dataset from the ‘22HP14’ experiment described above but by only using the counts made with the human eye. The hourly weather data were retrieved from a local weather station (INRAE #31035002; https://agroclim.inrae.fr/climatik/): temperature (actinothermic index at 50 cm, °C), relative humidity (%), photosynthetically active radiation (J/cm^2^) and wind speed (km/h). As described above, we also restricted the analyses to the hours of activity of each insect class, to decrease the number of ‘zeros’ and data overdispersion. All the statistical analyses ran to assess the differences of insect visitation frequency between sunflower genotypes are detailed in Supplementary data. All the statistical analyses were performed with the software R, version 4.2.3 (R Core team, 2023). The R code for analyses is available at https://forge.inrae.fr/astr/public/pollicrop_toolbox.

## 3. Results

### 3.1. Evaluation of the model: performance scores

After training and validation of the two versions of PolliCrop, one trained without and one with images lacking flower-visiting insects, both versions were tested with images of plants in the subsample of the image set not previously used for the training and validation (see section 2.3). The resulting performance scores are presented in Table 3. Regarding model precision, both PolliCrop versions hallucinated non-*Bombus* bees or bumble bees, on ca. 22% and 33% of images, respectively. However, PolliCrop2 hallucinated fewer lepidopterans (11% of images) than PolliCrop1 (22%). Regarding the recall, PolliCrop1 missed fewer non-*Bombus* bees, bumble bees and lepidopterans (16%, 10% and 9% of images, respectively) than PolliCrop2 (21%, 29% and 12%). As expected, PolliCrop2 decreased the recall, especially for bumble bees (-15%; -5% for non-*Bombus* bees and -4% for lepidopterans), but contrary to expectations it improved the precision only for lepidopterans (+11%), while the precision for non-*Bombus* bees and bumble bees was not impacted.

**Table 3.**
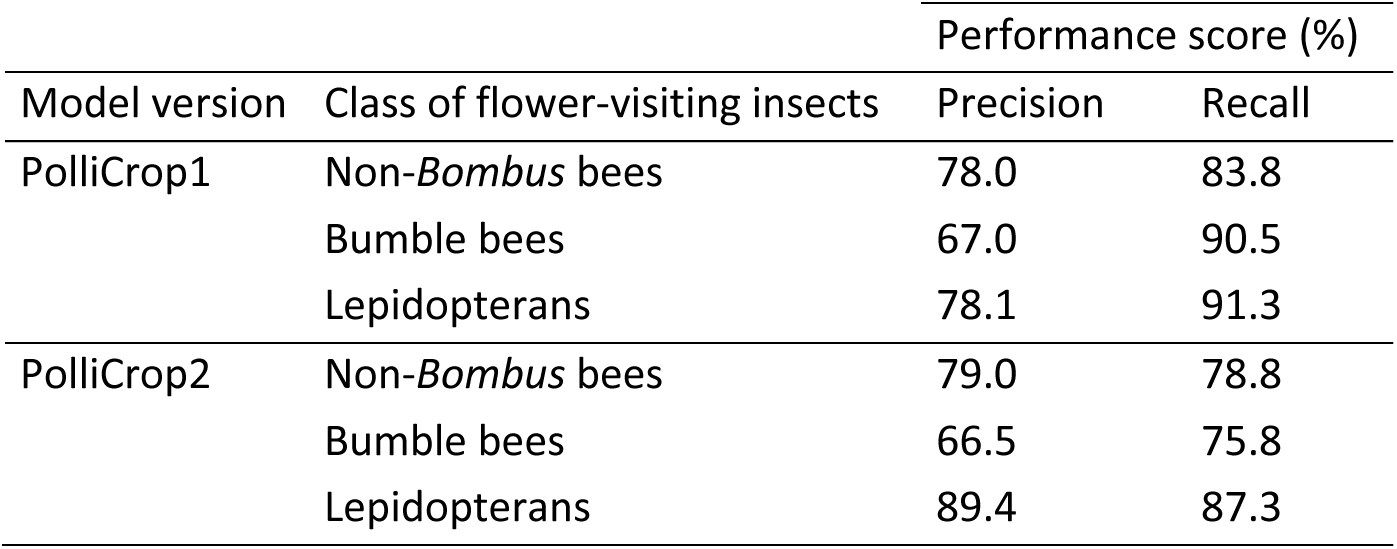
Performance scores calculated at the object level from the testing image sets of the two PolliCrop versions.

| Model version | Class of flower-visiting insects | Performance score (%) |  |
| --- | --- | --- | --- |
|  |  | Precision | Recall |
| PolliCrop1 | Non-*Bombus* bees | 78.0 | 83.8 |
|  | Bumble bees | 67.0 | 90.5 |
|  | Lepidopterans | 78.1 | 91.3 |
| PolliCrop2 | Non-*Bombus* bees | 79.0 | 78.8 |
|  | Bumble bees | 66.5 | 75.8 |
|  | Lepidopterans | 89.4 | 87.3 |

PolliCrop was then evaluated with two image sets (see section 2.4): the complete image set ‘22HP14’ which was used to train the model and composed of 8,970 images, and a sample of 1,024 images of the image set ‘24EV16’ that were not used to train the model. The two PolliCrop versions predicted the three flower-visiting insect classes well, between 96% and 100% of images, over the two image sets (accuracy; Tables 4 and 5). Regarding the precision, the two PolliCrop versions hallucinated minimal non-*Bombus* bees over the two image sets (< 10% of images for 22HP14; < 15% for 24EV16). PolliCrop1 hallucinated fewer bumble bees and lepidopterans (ca. 33% of images) than PolliCrop2 for 22HP14 (ca. 45%; Table 4), while PolliCrop2 hallucinated fewer bumble bees (17%) and lepidopterans (37%) than PolliCrop1 for 24EV16 (ca. 47% over the two classes; Table 5). Regarding the recall, PolliCrop1 missed quite a few insects, between 10% and 17% of images over the three classes and two image sets (except 0% for bumble bees for 24EV16, and 29% for lepidopterans for 22HP14), while PolliCrop2 missed more insects than PolliCrop1, between 24% and 49% of images over the three classes and two image sets (except 0% for bumble bees for 24EV16; Tables 4 and 5). As expected, PolliCrop2 decreased the recall, between 10% and 20% for the three insect classes and both image sets (except for bumble bees for 24EV16 for which the recall did not change). Contrary to expectations, it improved the precision only for bumble bees (+28%) and lepidopterans (+12%) for 24EV16, while it decreased the precision for bumble bees (-18%) and lepidopterans (-7%) for 22HP14. It also decreased the precision slightly for non-*Bombus* bees for both image sets (-3%).

**Table 4.** Evaluation at the image level of the two versions of PolliCrop on the image set ‘22HP14’, composed of 8,970 images.

| Model version | Class of flower-visiting insects | Performance score (%) |  |  |
| --- | --- | --- | --- | --- |
|  |  | Accuracy | Precision | Recall |
| PolliCrop1 | Non-*Bombus* bees | 98.5 | 95.2 | 86.1 |
|  | Bumble bees | 99.9 | 68.0 | 89.5 |
|  | Lepidopterans | 97.2 | 66.0 | 71.0 |
| PolliCrop2 | Non-*Bombus* bees | 97.4 | 92.2 | 75.8 |
|  | Bumble bees | 99.8 | 50.0 | 73.7 |
|  | Lepidopterans | 96.3 | 58.9 | 51.4 |

**Table 5.** Evaluation at the image level of the two versions of PolliCrop on a sample of 1,024 images of the image set ‘24EV16’.

| Model version | Class of flower-visiting insects | Performance score (%) |  |  |
| --- | --- | --- | --- | --- |
|  |  | Accuracy | Precision | Recall |
| PolliCrop1 | Non-*Bombus* bees | 98.0 | 89.3 | 84.8 |
|  | Bumble bees | 99.2 | 55.6 | 100 |
|  | Lepidopterans | 96.6 | 50.9 | 83.3 |
| PolliCrop2 | Non-*Bombus* bees | 97.1 | 85.9 | 72.3 |
|  | Bumble bees | 99.8 | 83.3 | 100 |
|  | Lepidopterans | 97.5 | 63.2 | 66.7 |

### 3.2. Assessment of the ability of the two PolliCrop versions to correctly predict the insect visitation frequency of three sunflower genotypes

Due to experimental constraints, the three genotypes did not benefit from the same level of monitoring. IR was the most monitored genotype, with a total of 417 monitoring hours over the 6 plants monitored (see Table 6 for the number of images), and a mean of 70 hours per plant. XRQ had a total of 258 monitoring hours over 6 plants, with a mean of 43 hours per plant. CI was the least monitored genotype, with 88 monitoring hours over 3 plants, and a mean of only 29 hours per plant. CI was especially undersampled during the night, with only 22 monitoring hours cumulative for the 3 plants from 20:00 to 08:00 (= the activity period of lepidopterans, see below). Overall, the number of visitations counted with the eye on the images amounted to 808 non-*Bombus* bees, 435 lepidopterans, and only 19 bumble bees (Table 6).

**Table 6.** Number of insects visiting the sunflower heads counted with the human eye on the images of the set ‘22HP14’.

| Class of flower-visiting insects | Genotype |  |  |  |
| --- | --- | --- | --- | --- |
|  | XRQ | IR | CI | Total |
| Non-*Bombus* bees | 304 | 445 | 59 | 808 |
| Bumble bees | 7 | 10 | 2 | 19 |
| Lepidopterans | 117 | 312 | 6 | 435 |
| *Total* | *428* | *767* | *67* | *1,26* |
| Total number of images | 3,048 | 4,948 | 974 | 8,970 |
| Number of images from 08:00 to 21:00 (day) | 1,719 | 2,707 | 805 | 5,231 |
| Number of images from 20:00 to 08:00 (night) | 1,469 | 2,445 | 203 | 4,117 |

Manual counts made with the eye enabled us to assess the insect activity during day and night (Fig. 3). Non-*Bombus* bees started to visit the sunflower heads from 08:00, shortly after sunrise, until 21:00 soon after sunset. The activity on XRQ and IR was quite similar, with a progressive increase in the early morning and a progressive decline in the late afternoon, with a slightly variable plateau around 0.25 bees per image, between 10:00 and 18:00. The activity on CI was noticeably lower, especially in the morning and the late afternoon. Bumble activity was comparatively near zero on the three genotypes, occurring only in the afternoon, from 12:00 to 20:00. Lepidopteran activity started from 20:00 at sunset and continued until 08:00 at sunrise. The activity of lepidopterans during the daylight hours were almost null. Lepidopterans showed similar activity on the three genotypes, with a more intense activity in the early night, sharply declining after 02:00. In the following results, the analysis of activity was limited to the day period, from 08:00 to 21:00, for both classes of non-*Bombus* bees and bumble bees, and to the night period, from 20:00 to 08:00, for lepidopterans.

**Fig. 3.**
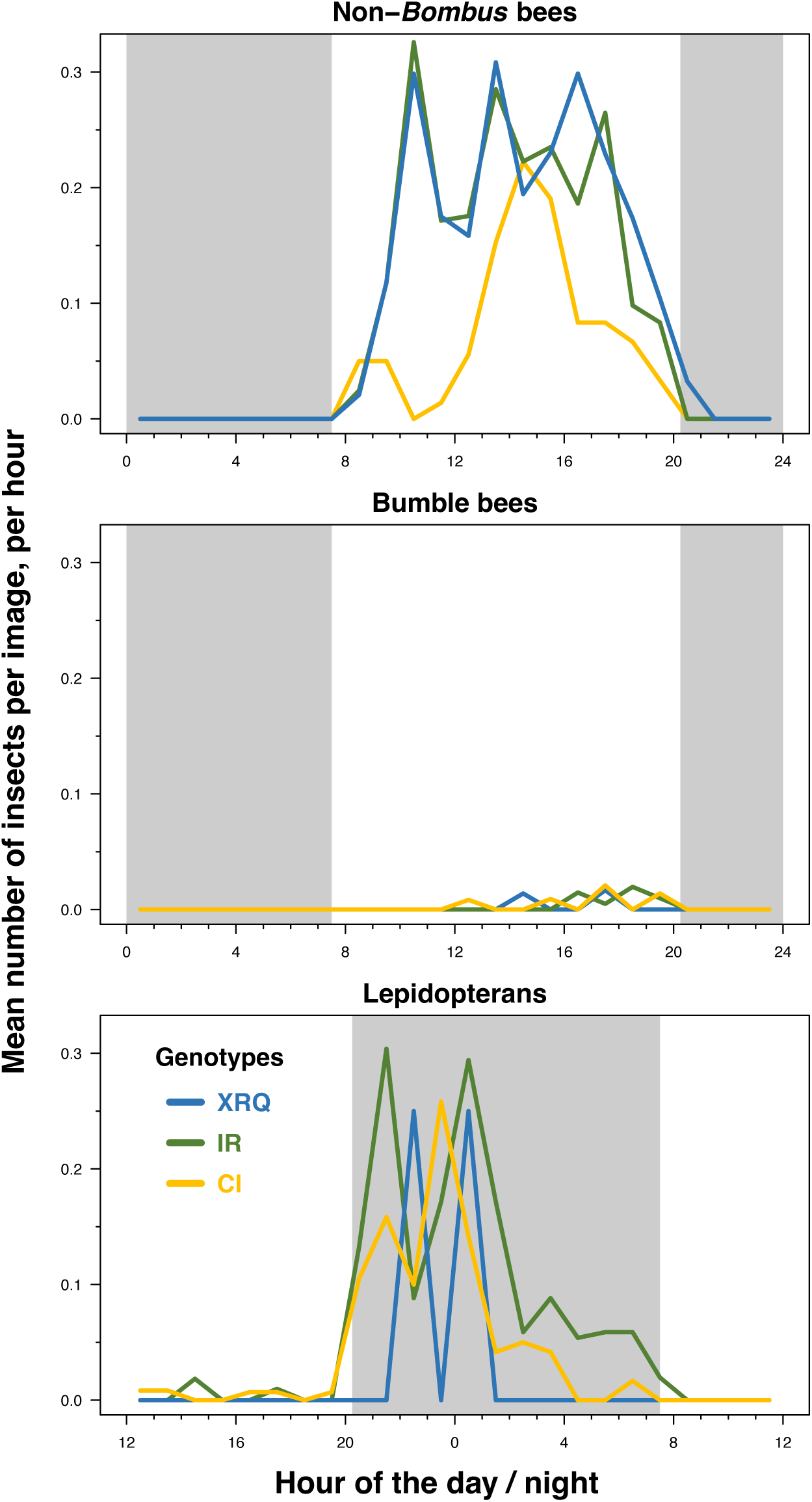
Number of insects visiting the sunflower heads during the daylight hours (light area) and the night period (grey area). Time is expressed in local time: e.g., 02:00 corresponds to midnight (00:00 UTC). Numbers are derived from the counts made with the human eye. Approximately 12 images (sometimes a bit less) were captured and summed per hour each day, per plant.

When comparing the counts of insects made with the eye and the two versions of PolliCrop, PolliCrop1 performed the best, successfully predicting the mean number of non-*Bombus* bees and lepidopterans per image on XRQ and IR in an error range of ±10% (Fig. 4). PolliCrop2 performed worse, underestimating non-*Bombus* bees by 20% on IR, and lepidopterans by 18% and 36% respectively on IR and XRQ. Both PolliCrop versions underestimated the mean number of bumble bees per image on IR, by -20% for PolliCrop1 and -10% for PolliCrop2, while they overestimated them on XRQ, by +57% and +139% respectively for PolliCrop1 and PolliCrop2. On the other hand, both versions performed poorly for the three insect classes on CI, underestimating non-*Bombus* bees by -53% and -66% respectively for PolliCrop1 and PolliCrop2, overestimating bumble bees and lepidopterans by +200% and +83% respectively for PolliCrop1, and underestimating bumble bees and lepidopterans by -50% and -100% respectively for PolliCrop2. Overestimations of insect visitations occur when the precision of PolliCrop for the given insect class and genotype is low (< 70%) while the recall is correct (> 80%), or when both scores are low (< 70%) but precision is even lower than recall (Table 7). Conversely, underestimations occur when the recall is low (< 70%) while the precision is correct (> 80%), or when both scores are low (< 70%) but recall is even lower than precision (Table 7). It can occur sometimes that both scores are quite low, but not too low (between 60% and 80%), and compensate for each other, resulting in an over- or underestimation of < 10% (e.g., prediction of PolliCrop1 for lepidopterans on XRQ and IR; Table 7; Fig. 4).

**Fig. 4.**
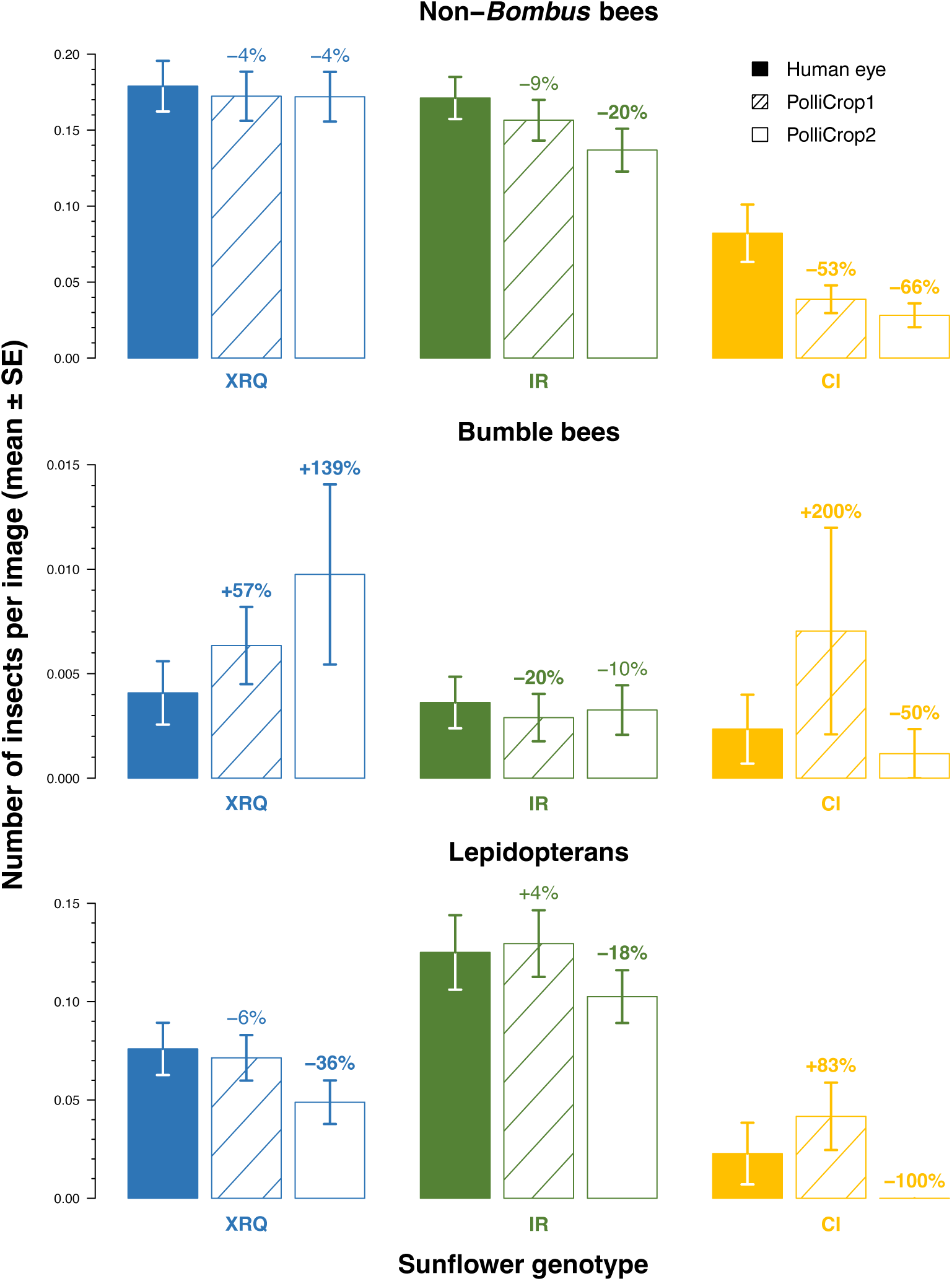
Insect activity on the sunflower heads assessed from the human eye and the predictions of the two versions of PolliCrop, in relation with the genotype and the insect class. The counts were limited to the activity period of each insect class, during the day period from 08:00 to 21:00 for both classes of non-*Bombus* bees and bumble bees, and to the night period from 20:00 to 08:00 for lepidopterans. Numbers above the bars are the % of increase or decrease of the prediction of PolliCrop in comparison with the counts made with the human eye. The % > |±10%| are highlighted in bold.

**Table 7.** Performance scores calculated at the image level for the two versions of PolliCrop run on the image set ‘22HP14’, according to the genotype.

|  |  | Genotype |  |  |  |  |  |
| --- | --- | --- | --- | --- | --- | --- | --- |
|  |  | XRQ | IR | CI | XRQ | IR | CI |
| Model version | Class of flower-visiting insects | Precision (%) |  |  | Recall (%) |  |  |
| PolliCrop1 | Non-*Bombus* bees | 96.8 | 94.2 | 94.1 | 93.4 | 85.6 | 54.2 |
|  | Bumble bees | 63.6 | 100 | 33.3 | 100 | 80 | 100 |
|  | Lepidopterans | 79.4 | 63.4 | 15.4 | 73.9 | 70.5 | 33.3 |
| PolliCrop2 | Non-*Bombus* bees | 96.4 | 88.4 | 95.7 | 91.7 | 70.3 | 37.3 |
|  | Bumble bees | 33.3 | 77.8 | 100 | 85.7 | 70.0 | 50.0 |
|  | Lepidopterans | 71.2 | 56.9 | 100 | 45.6 | 55.0 | 0 |

### 3.3. Comparison of statistical models to assess the differences of insect visitation frequency between sunflower genotypes

As described in the previous section, the analysis was limited to the activity period of each insect class, during the day from 08:00 to 21:00 for both classes of non-*Bombus* bees and bumble bees, and the night from 20:00 to 08:00 for lepidopterans, with most of the activity occurring during the first half of the night, between 20:00 and 02:00. After top-down model selection to choose the optimal random effect structures of GL(M)Ms to explain insect visitation frequency, the date effect was kept in the GLMMs without the weather variables for the three insect classes, but not in the GL(M)Ms with the weather variables except for lepidopterans (Tables 8 and S1). On the other hand, the plant effect was kept only in the GLMMs explaining the visitation frequency of non-*Bombus* bees, with or without the weather variables, and in the GLMM with the weather variables explaining the visitation frequency of lepidopterans.

During the recordings, photosynthetically active radiation ranged from 0 to 130 J/cm^2^ during the day between 08:00 and 21:00 (or from 0 to 26 J/cm^2^ during the night between 20:00 and 08:00), temperature from 16° to 37°C (11-26°C during the night), relative humidity from 23 to 97% (39-99%) and wind speed from 0 to 23 km/h (0-18 km/h). Three variables displayed pairwise |r| > 0.7: temperature with radiation (r = 0.78), and temperature with relative humidity (r = -0.78; Fig. S6). After selection by AIC, all the weather predictors were present in at least one model in the sets of most parsimonious models for the three insect classes (Table S2). The set of most parsimonious models included three models for non-*Bombus* bees: the complete model that included all the weather predictors and the models that included or excluded radiation or wind speed. This set included six models for bumble bees: these models included or excluded radiation, wind speed, or temperature with a squared term. And it included only two models for lepidopterans: the complete model and the model without wind speed. The three sets of most parsimonious models included temperature with a simple term and relative humidity in all models. The complete models with all the weather predictors were therefore chosen for comparisons with the models without the weather variables for all three insect classes.

Both models with or without the weather variables yielded similar results in terms of statistical significance, with a *P*-value threshold of 0.05; the genotype CI received fewer non-*Bombus* bees and lepidopterans than XRQ and IR, while XRQ and IR received similar visitation frequencies for both insect classes (Table 8; Fig. 5). Regarding bumble bees, the three genotypes received similar visitation frequencies. However, in terms of effect sizes, the two models yielded substantially different results. The model with the weather variables resulted in greater differences of non-*Bombus* bee visitation frequencies for IR and CI in comparison with XRQ than the model without the weather variables (Fig. 5). The bumble bee visitation frequencies were higher for IR and CI compared to XRQ with the model with the weather variables, while they were lower with the model without the weather variables. For lepidopterans, the two models yielded similar differences between IR, CI, and XRQ.

**Fig. 5.**
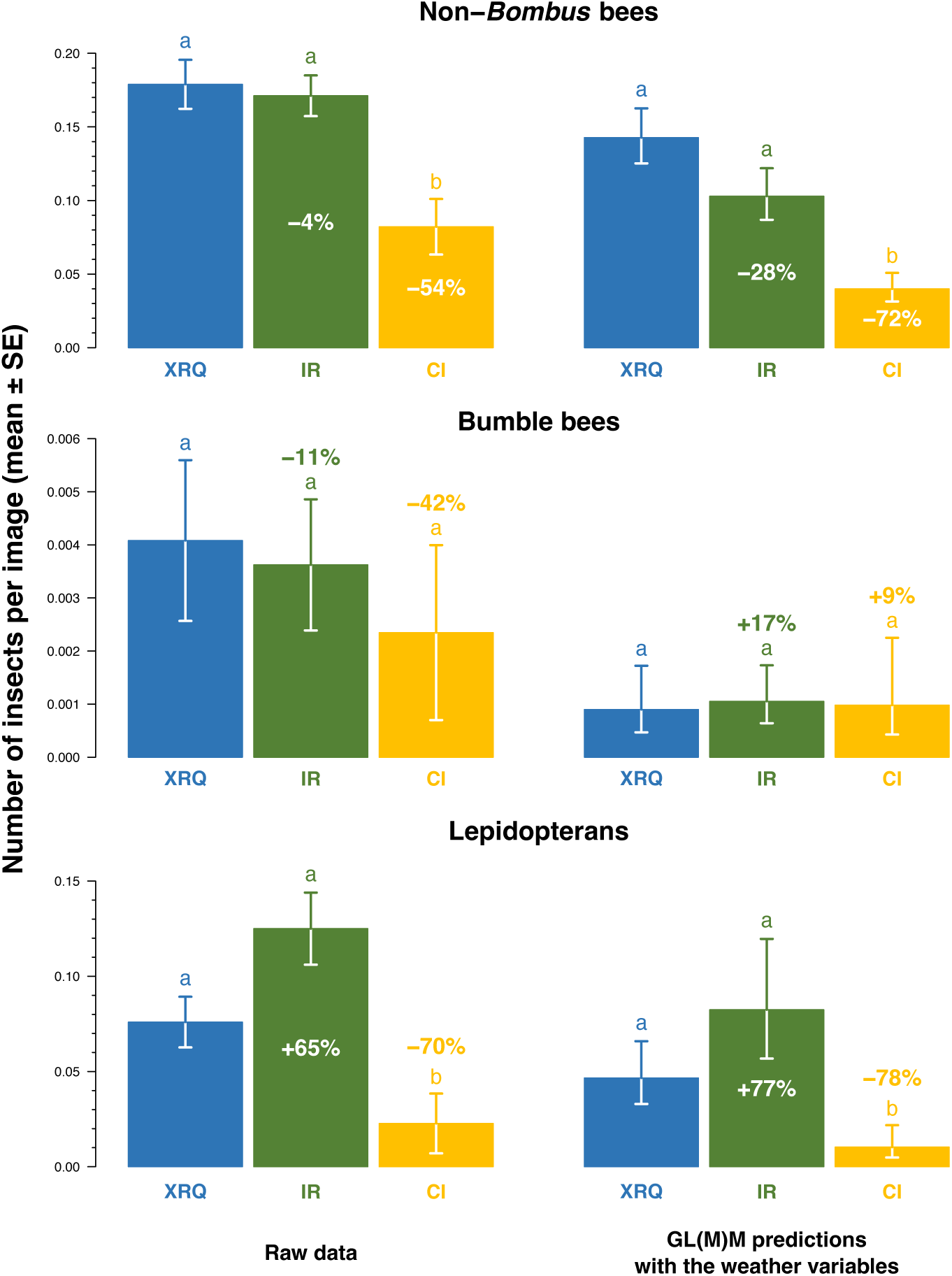
Insect activity on the sunflower heads assessed from the human eye, in relation with the genotype and the insect class. On the left, data are the averages directly calculated from the raw data. On the right, data are the predictions given by the complete GL(M)Ms with all the weather variables (Table 8). The counts were limited to the activity period of each insect class, during the day period from 08:00 to 21:00 for both classes of non-*Bombus* bees and bumble bees, and to the night period from 20:00 to 08:00 for lepidopterans. Percentages are the increases or decreases of IR and CI in comparison with XRQ. A different letter above the bars indicates a difference with *P* < 0.05. The statistics on the left are from the GLMMs without the weather variables, and from the complete GL(M)Ms with all the weather variables on the right (Table 8). The weather variables were all centered and standardized to estimate insect visitation frequencies per genotype on average weather values (see caption in Table 8 for the values).

**Table 8.**
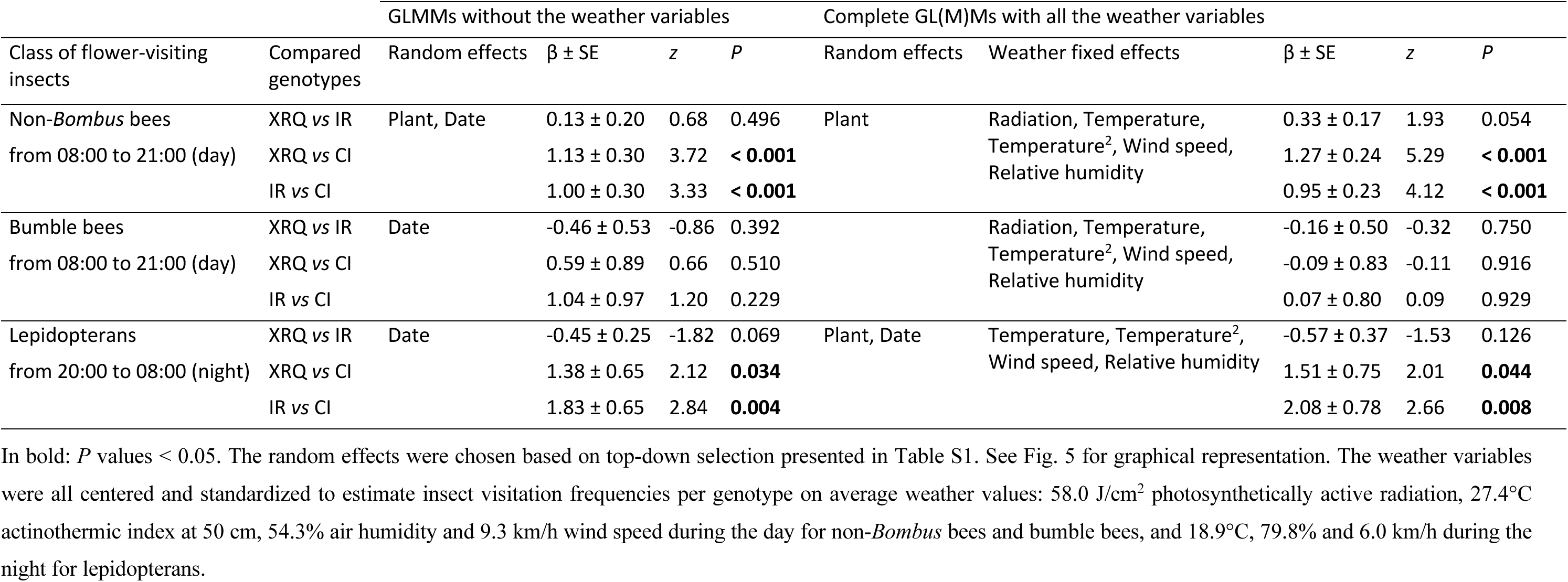
Pairwise comparisons of the number of insects per image between the three sunflower genotypes, estimated from the GL(M)Ms with or without the weather variables.

|  |  | GLMMs without the weather variables |  |  |  | Complete GL(M)Ms with all the weather variables |  |  |  |  |
| --- | --- | --- | --- | --- | --- | --- | --- | --- | --- | --- |
| Class of flower-visiting insects | Compared genotypes | Random effects | $\beta \pm SE$ | $z$ | $P$ | Random effects | Weather fixed effects | $\beta \pm SE$ | $z$ | $P$ |
| Non-*Bombus* bees<br>from 08:00 to 21:00 (day) | XRQ vs IR | Plant, Date | 0.13 ± 0.20 | 0.68 | 0.496 | Plant | Radiation, Temperature,<br>Temperature<sup>2</sup>, Wind speed,<br>Relative humidity | 0.33 ± 0.17 | 1.93 | 0.054 |
| | XRQ vs CI | | 1.13 ± 0.30 | 3.72 | **< 0.001** | | | 1.27 ± 0.24 | 5.29 | **< 0.001** |
| | IR vs CI | | 1.00 ± 0.30 | 3.33 | **< 0.001** | | | 0.95 ± 0.23 | 4.12 | **< 0.001** |
| Bumble bees<br>from 08:00 to 21:00 (day) | XRQ vs IR | Date | -0.46 ± 0.53 | -0.86 | 0.392 | | Radiation, Temperature,<br>Temperature<sup>2</sup>, Wind speed,<br>Relative humidity | -0.16 ± 0.50 | -0.32 | 0.750 |
| | XRQ vs CI | | 0.59 ± 0.89 | 0.66 | 0.510 | | | -0.09 ± 0.83 | -0.11 | 0.916 |
| | IR vs CI | | 1.04 ± 0.97 | 1.20 | 0.229 | | | 0.07 ± 0.80 | 0.09 | 0.929 |
| Lepidopterans<br>from 20:00 to 08:00 (night) | XRQ vs IR | Date | -0.45 ± 0.25 | -1.82 | 0.069 | Plant, Date | Temperature, Temperature<sup>2</sup>,<br>Wind speed, Relative humidity | -0.57 ± 0.37 | -1.53 | 0.126 |
| | XRQ vs CI | | 1.38 ± 0.65 | 2.12 | **0.034** | | | 1.51 ± 0.75 | 2.01 | **0.044** |
| | IR vs CI | | 1.83 ± 0.65 | 2.84 | **0.004** | | | 2.08 ± 0.78 | 2.66 | **0.008** |

Based on the complete model with all the weather variables and the multimodel averaging, non-*Bombus* bees and lepidopteran activity on sunflower heads responded to the temperature with a bell shape, with an optimum temperature of 30.1°C for non-*Bombus* bees and of 20.0°C for lepidopterans (*P* < 0.05 and RI = 100% for the squared term; Table 9). On the other hand, bumble bee activity decreased with a constant negative effect with the temperature (*P* < 0.05 and RI = 100% for the simple term). The activity of the three insect classes decreased with the relative humidity (*P* < 0.05; RI = 100%), while radiation and wind speed both did not show evidence of effect on these activities (*P* > 0.1; RI ≤ 51% for radiation and ≤ 45% for wind speed), but with weak evidence for a negative effect of wind speed on non-*Bombus* bee activity (*P* < 0.1; RI = 79%). Overall, based on the multimodel averaging, the effect size was the greatest for the temperature for non-*Bombus* bee activity, while the effect size of the relative humidity was the greatest for bumble bee activity, and while the effect sizes of both temperature and relative humidity were quite similar for lepidopteran activity (Table 9).

**Table 9.** Predictions and statistics of the weather predictors, estimated from the GL(M)Ms with the weather variables.

| Class of flower-visiting insects | Random effects | Weather predictor | Complete model with all the weather variables |  |  | Multimodel averaging |  |
| --- | --- | --- | --- | --- | --- | --- | --- |
| | | | $\beta \pm SE$ | $z$ | $P$ | Relative importance (RI) | $\beta \pm SE$ |
| Non-*Bombus* bees<br>from 08:00 to 21:00 (day) | Plant | Radiation | 1.72 ± 1.71 | 1.01 | 0.313 | 51% | 2.18 ± 1.66 |
| | | Temperature | 14.45 ± 3.33 | 4.34 | **< 0.001** | 100% | 15.00 ± 2.88 |
| | | Temperature<sup>2</sup> | -12.56 ± 1.83 | -6.85 | **< 0.001** | 100% | -12.60 ± 1.80 |
| | | Wind speed | -2.63 ± 1.57 | -1.67 | 0.094 | 79% | -3.01 ± 1.49 |
| | | Relative humidity | -8.13 ± 2.21 | -3.68 | **< 0.001** | 100% | -8.20 ± 2.10 |
| Bumble bees<br>from 08:00 to 21:00 (day) | | Radiation | 5.30 ± 9.66 | 0.55 | 0.583 | 24% | 6.77 ± 9.01 |
| | | Temperature | -41.59 ± 16.22 | -2.56 | **0.010** | 100% | -36.95 ± 9.68 |
| | | Temperature<sup>2</sup> | -11.90 ± 8.26 | -1.44 | 0.150 | 48% | -9.38 ± 7.62 |
| | | Wind speed | -8.10 ± 8.46 | -0.96 | 0.338 | 28% | -7.31 ± 7.76 |
| | | Relative humidity | -48.72 ± 14.90 | -3.27 | **0.001** | 100% | -48.12 ± 12.67 |
| Lepidopterans<br>from 20:00 to 08:00 (night) | Plant, Date | Temperature | 6.83 ± 4.80 | 1.42 | 0.155 | 100% | 4.68 ± 3.91 |
| | | Temperature<sup>2</sup> | -7.78 ± 3.05 | -2.55 | **0.011** | 100% | -8.24 ± 2.87 |
| | | Wind speed | -5.87 ± 4.61 | -1.27 | 0.203 | 45% | -5.87 ± 4.61 |
| | | Relative humidity | -8.72 ± 3.16 | -2.76 | **0.006** | 100% | -8.31 ± 2.88 |

## 4. Discussion

### 4.1. Performance of PolliCrop to predict the insect classes at the image level

Overall, the PolliCrop deep learning model was efficient at successfully predicting most of images, between 96% and 100%. This ability is largely driven by the high rate of images without insects (> 85% in the two examples of image sets), mostly correctly predicted by the model. The two versions of PolliCrop were also efficient at predicting non-*Bombus* bees, with a few “hallucinations” for both versions, but more bees were missed by PolliCrop2. PolliCrop2 hallucinated fewer bumble bees and lepidopterans than PolliCrop1, although it depended on the image set. Learning “empty” images, i.e., without insect, did not necessarily improve model’s precision. On the other hand, PolliCrop1 missed fewer insects (non-*Bombus* bees, bumble bees, and lepidopterans) than PolliCrop2 on all image sets. Thus, PolliCrop1 is recommended against PolliCrop2, although it should depend on the context and image set. If PolliCrop2 yields fewer “hallucinations” on a given image sample of an image set, and that the image set includes a high rate of “empty” images or of poor quality (e.g., blurred images or images underexposed to light), then running PolliCrop2 may be preferred. On the other hand, if one would like to check the images after model running in an image set with a high rate of “empty” images, running a version with a low rate of missed insects (but a high rate of “hallucinations”) would be preferred against a version with a low rate of hallucinations (but a high rate of missed insects), since reviewing the few images with an insect predicted by the model (= ‘positive’ images) would take less time than reviewing all images for which the model predicted insect absence. Regardless, the model was effective at predicting insects on new image sets that were not used to train the model. However, to confirm if the model is well generalizable to new image sets of sunflower heads, captured in other contexts, countries, seasons, insect communities, etc., the model should be tested with these new types of image sets. Most likely, images of these new sets will need to be segmented and identified at the insect morphogroup level to increase the model’s generalizability. The “new” image set tested here came from the exact same location, so not an evident proof of generalizability. Future improvements of the model should also aim to increase the performance scores per insect class, the “intra-class” detection, e.g., between bumble bee morphogroups (white-end *Bombus*, red-end *Bombus*, brown *Bombus*; e.g., Bjerge et al., 2023a,c; Smith et al., 2026), the number of classes detected (e.g., hoverflies, Vespidae, *Xylocopa*), as well as other sunflower head morphs and other crop species (e.g., apple crop; Smith et al., 2026). The model will therefore gain from increasing the number of images segmented, especially with the insect morphogroups poorly represented in the current image set used to train the model, and increasing the number of crop species monitored with cameras, e.g., by pooling all the images available in the different projects of deep learning model developments for flower-visiting insect detection (Spiesman et al., 2021; Bjerge et al., 2023a,b,c, 2024; Stark et al., 2023; Sittinger et al., 2024; Alex et al., 2025; Serra-Marin et al., 2025; Ştefan et al., 2025; Bernauer et al., 2026; Gardiner et al., 2026; Smith et al., 2026).

### 4.2. Performance of PolliCrop to predict the insect visitation frequency at the level of sunflower genotype

When comparing the counts of insect visitations made on sunflower heads by the human eye with those predicted by PolliCrop on a complete image set that was used to train the model, the version PolliCrop1 performed the best for two genotypes out of three, XRQ and IR. In particular, PolliCrop1 predicted well the visitation frequency of non-*Bombus* bees and lepidopterans per image in a range of ±10%. As expected, when the scores of precision and recall are equivalent and not too low, by aggregating multiple images and under the assumption of mutually independent image captures, these two scores can compensate for each other, resulting in an over- or underestimation of insect visitation frequency < 10%, as is the case of the predictions of lepidopterans by PolliCrop1. The estimations and additions of these two performance scores for a given insect class can therefore inform on the level of over- or underestimation of insect visitation frequency by the model. The assumption of mutually independent image captures is not met by the nature of ‘time series’ of image captures. But this limitation can be reduced by increasing the number of plants monitored.

It is also interesting to note that the performance of the model can depend on the sunflower genotype. Both PolliCrop versions were, for instance, poor at predicting the insect visitation frequency for the genotype CI for the three insect classes. Several factors may explain this poor performance. First, this outcome can be explained by the low level of monitoring of this genotype, both in terms of sample size and hours monitored per plant, resulting in quite small total numbers of insect visitations recorded, especially for bumble bees and lepidopterans, likely increasing the error percentage of prediction as a consequence. The poor prediction of PolliCrop for CI may also be explained by the unique head morphology of CI, featuring a darker general appearance and more prominent stigmas compared to a “standard” cultivated sunflower head (see Fig. S5). These distinctive features may explain the increased rate of missed non-*Bombus* bees by the model, through an increased confusion of their morphology with the background. A third explanation for the low performance of PolliCrop for predicting the insect visitation frequency on CI may be the elevated frequency of blurred images observed during visual review. Across all three genotypes, the poor performance of PolliCrop for predicting the visitation frequency of bumble bees may be explained by the overall very low frequency of bumble bee visitations. The very small total raw number of bumble bee visitations recorded may then again inevitably increase the error percentage of prediction.

Our recommendations for reaching the highest level of performance as possible for PolliCrop to predict the insect visitation frequency in a given image set would be (*i*) to monitor a sufficient number of plants per genotype; (*ii*) to monitor the plants during a sufficient period of time, ideally during the whole blooming period of the heads; (*iii*) to maximize the quality of the images, for instance by avoiding as many as possible blurred images. For the future, the general performance of PolliCrop to predict insect visitations in new images sets with a variety of sunflower morphs could be increased by increasing the size of the training image set with a variety of sunflower morphs, including branched and single headed genotypes, and by retraining the model with this augmented image set. Regardless, it can be useful to estimate performance scores per genotype, by reviewing with the eye an equal number of sampled images.

### 4.3. Circadian clock and effect of weather conditions on insect foraging

The daily pattern of activity of bees found in our results is consistent with what has been reported in the literature for foraging activity recorded during summer: sharp increase and decrease in the morning and evening, respectively, with a plateau during the day (Stelzer and Chittka, 2010; Steen, 2017). This daily pattern is not driven by the absolute levels of light intensity, but by the circadian clock which is entrained by the natural daily cycles in light intensity and the associated temperature fluctuations (Stelzer and Chittka, 2010; Steen, 2017). This is how bumble bees showed a foraging activity with a day/night cycle even with permanent daylight conditions during summer north of the Arctic circle (Stelzer and Chittka, 2010). This can explain why variations of radiation during the day did not show evidence of effect on bee visitations to sunflowers in our results.

Regarding the temperature effect, while it is theoretically well established that the foraging activity of bees should decrease beyond an optimum temperature (Colinet et al., 2015), empirical studies are scarce in the literature. Our value of 30°C found for the optimum activity of non-*Bombus* bees on sunflower heads can for instance be compared with the optimum of 33°-37°C found for honey bees foraging on strawberry flowers in Australia (Jaboor et al., 2022), or of 24°C for honey bees foraging on raspberry flowers late Spring in Norway (Nielsen et al., 2017). This optimum may probably depend on the season, crop visited and environmental context, but further studies are required. A negative effect of the temperature was found on bumble bee activity, but this probably arises from the very low number of bumble bees recorded, resulting in poor statistical power. Kenna et al. (2021) found an optimum temperature of 25°C for the flight performance of bumble bees, while in our study the day temperatures recorded covered a range above and below this optimum. The negative effect of relative humidity found on non-*Bombus* bee and bumble bee activity is consistent with the observations of Sanderson et al. (2015) of bumble bee activity at nest entrance. Sanderson et al. (2015) suggested that this effect could be due to the decrease of nectar concentration in sugars with increasing relative humidity (Pacini and Nepi, 2007), resulting in a decrease of its nutritional value per unit volume. But this hypothetical mechanism needs empirical experiments to be confirmed. This effect could also be an artifact resulting from the strong negative correlation between temperature and relative humidity, or from a potential positive correlation between relative humidity and rainfall. The effect found for relative humidity could be indirect resulting only from the combined effects of temperature and rainfall. The weak negative effect of wind speed found on non-*Bombus* bee activity is consistent with what has been previously reported in the literature with the foraging activity of honey bees, bumble bees and other wild bees (Pinzauti, 1986; Vicens and Bosch, 2000; Sanderson et al., 2015; Ngo et al., 2021); although Pinzauti (1986) found a sharp decrease of honey bee foraging until 0 between 10 and 16 km/h. The activity period observed for lepidopterans, occurring mostly during the first half of the night (between 20:00 and 02:00 in our study), is consistent with the observations made by Knop et al. (2018) on night lepidopterans visiting meadow flowers (mostly between 22:00 and 02:00). For our observations, this activity period may be the result of nectar secretion that occurs only during the day for sunflower (Stan Chabert, unpublished data). Moths would visit sunflower heads primarily during the first half of the night to feed on residual nectar left by bees during the day, and would leave the heads when all this residual nectar has been completely consumed. An optimum of 20°C was found for lepidopteran activity in our study, while Knop et al. (2018) found only a simple positive effect of temperature.

Finally, including or excluding the weather variables in the statistical model did not change the statistical conclusions made on the differences of visitation frequencies between the three sunflower genotypes for the three insect classes. This observed insensitivity to weather conditions is probably due to the relative short period of ten days monitoring insect activity. During such a short period, most of the plants of the three genotypes finally bloomed at the same time. Including the weather variables in the model would be especially relevant for comparing genotypes with non-overlapping bloom and with strong variability of weather conditions during the whole monitoring period. In the future, this type of analysis should directly include general knowledge about the flower-visiting insect responses to the weather conditions via priors in a Bayesian approach (Clark and Gelfand, 2006; Korner-Nievergelt et al., 2015). This approach would make it possible to overcome the specific weather conditions inherent to each experiment. But this requires first to exhaustively investigate the insect activity on flowers in relation with the weather conditions over different seasons. For instance, the recently developed devices to automatically count the honey bees entering and leaving the hive per unit of time could be used for that (Meikle and Holst, 2015; Clarke and Robert, 2018; Odemer, 2022).

### 4.4. Community of insect pollinators visiting sunflower heads

The high frequency of honey bee visitations to sunflower heads found in our image set is consistent with the large dominance of honey bees observed worldwide in the literature for the insect pollinators visiting sunflower, in countries other than North America (see references in Brown and Cunningham, 2019; Chabert et al., 2022; Aubouin et al., 2026). Sunflower is native to North America (Kantar et al., 2015), and in this region, heads receive more visits from wild bees, with a higher species diversity observed than anywhere else, than honey bees (see references in Brown and Cunningham, 2019; Chabert et al., 2022). In France for instance, only 11 wild bee species, including bumble bees, have been recorded at least 5 times in the databases of Rollin et al. (2015) and Aubouin et al. (2026). For the future development of PolliCrop, to extend the ability of the model to identify the insects visiting sunflower in all countries, the training image set should be reinforced with images coming from a variety of countries, and especially from North America.

Moths were surprisingly very abundant in our image sets, present in 30% of the images that included at least one “pollinator” on the head (Table 1). They foraged on sunflower mostly during the night (see above), but also a little during the day (Fig. 3). Their visitation frequency to sunflower heads almost reached that of the non-*Bombus* bees: 76% of the non-*Bombus* bee frequency on IR and 44% on XRQ (Fig. 5). Carvalheiro et al. (2011) also recorded moths visiting sunflower heads in fields during the day in South Africa, while Torretta et al. (2009) observed a high moth abundance during the night in Argentina over 3 years. In this last study, the richness of moth community was unexpectedly high, including 3 families, 10 subfamilies, 24 genus and 34 species. This study contributes therefore to the growing literature showing that nocturnal insects represent a substantial fraction of the flower-visiting insects, overlooked so far, potentially contributing to pollination (Macgregor and Scott-Brown, 2020; Requier et al., 2023). In the case of sunflower, since pollen is emitted in the early morning (Creux et al., 2021), moths visiting sunflower at night most probably make limited contributions to pollination (but see Nderitu et al., 2008).

### 4.5. Experimental design and interpretation of attractiveness

Caution should be paid to the experimental design for interpretation of plant attractiveness. (*i*) Honey bees and bumble bees usually forage following a linear pattern within the same rows of plants (Free and Spencer-Booth, 1964; Cresswell et al., 1995; Jackson, 1996; Brittain et al., 2013; Mallinger et al., 2024). Mixing plant genotypes within rows would therefore reinforce the experimental design for testing the preference of bees towards certain genotypes. (*ii*) Insect pollinators learn to associate a set of floral traits (morphology, color, volatile organic compounds, electric field, etc.) characterizing a plant’s genotype to its level of resources offered (Wright and Schiestl, 2009; Frachon et al., 2021). There is therefore probably a minimum “mass” of plants of each genotype to provide within the experimental plots, to give the insects enough plants to learn which genotypes are the most profitable. (*iii*) This is not because a plant genotype is weakly attractive to insect pollinators in experimental plots mixing multiple genotypes, that this genotype will also be weakly visited by insect pollinators in commercial fields offering only this single genotype. The level of insect visitation to a field depends on the level of resources offered by both the field and the environment in the surroundings. (*iv*) Honey bee and bumble bee colonies can be added to the experimental plots to ensure the presence of both these taxa, bearing in mind that the presence of one taxon can influence the distribution on genotypes of the other taxa (Ferguson et al., 2021).

### 4.6. Conclusion

In this study, we developed PolliCrop, a deep learning model that automatically detects and identifies three classes of the main insect pollinators from images of sunflower heads captured during the day and the night with satisfying performance scores. This model will be particularly useful for high-throughput phenotyping of the attractiveness of plant genotypes to flower-visiting insects. The model was developed first on sunflower, a plant with all the flowers concentrated in a single inflorescence, making this species particularly well adapted for monitoring flower-visiting insects at the plant level with a camera. PolliCrop can be extended in the future to other crop species by adding new segmented images captured on these crops to the training image set. An important improvement for the overall pipeline of high-throughput phenotyping of attractiveness of plants to flower-visiting insects will be to develop an embedded system to directly detect, identify, and record insects on video monitoring and by saving only the count data on-device. This capacity will enable the monitoring of the whole blooming period of plants in a continuous way, yielding the exact number of insect pollinators that visited a set of flowers during the whole life of these flowers. This improved tool could have multiple applications, including studying the relationship between the total number of pollinator visitations received by plants and theirs associated pollination outcomes, or the foraging behavior of pollinators, e.g., if they collect pollen *vs.* nectar, their time spent per plant, the possible interference between conspecifics *vs.* heterospecifics etc.

## Author contributions

NBL, JBS and BKB conceived the study. NBL and SE acquired the fundings and oversaw the study. NB, OC and MG collected the images. SC, SE, OC and SH identified the insect classes. OC, SC, NB, CD, MG, SH, NBL, GT and SE reviewed and segmented the images. JBS and RG developed the deep learning model. JBS, NB and NBL prepared the Python files and routines. SC and GT analyzed the data. SC wrote the first draft and created the figures and tables. BKB, SH, NL, SE, JBS and GT reviewed and edited the manuscript. All authors approved the manuscript in its final form.

## Funding

This work is part of HelEx and AGRI4POL projects funded by the European Union’s Horizon Europe Research and Innovation Actions programme under grant agreements #101081974 and #101181146. This research used the PHENOME-EMPHASIS facility Phenotoul-Heliaphen (Phenome-ANR-11-INBS-0012; https://doi.org/10.15454/1.5483266728434124E12), was part of the French Laboratory of Excellence project “TULIP” (ANR-10-LABX-41; ANR-11-IDEX-0002-02) and of the project Heliopollen funded by Promosol, and was supported by the France-Berkeley Fund. This work is part of the project Attracthol granted by Plant2Pro^®^ Carnot Institute in the frame of its 2022 call for projects. Plant2Pro^®^ is supported by ANR (agreement #22-CARN-024-01 – 2021).

## Declaration of competing interests

The authors declare no competing interests.

## Acknowledgments

The authors are grateful to Sarah Lawniczak, Flavie Brunet, Edith Titaud, Julien Marc, Marie-Sophie Metzdorf, Juliette Blanc, Jade Boisbouvier and Louka Le Sager who helped in reviewing and segmenting the images and/or collecting the images.

## Supplementary data

MMC S1. Supplementary Material includes additional figures and tables as well as additional information for the material and method section.

## Data availability

Data, spreadsheets, Python files and R code are available at: https://forge.inrae.fr/astr/public/pollicrop_toolbox.

## Notes

https://forge.inrae.fr/astr/public/pollicrop

https://forge.inrae.fr/astr/public/pollicrop_toolbox

### Competing Interest Statement

The authors have declared no competing interest.

### Summary of Updates

Several points in the main manuscript and in the Supplementary materials have been updated. In particular, the URL quoted in the mnuacript have been created and updated.

## References

Alex, A. J., Barnes, C. M., Machado, P., Ihianle, I., Markó, G., Bencsik, M., & Bird, J. J. (2025). Enhancing pollinator conservation: Monitoring of bees through object recognition. Computers and Electronics in Agriculture, 228, 109665.

Alison, J., Alexander, J. M., Diaz Zeugin, N., Dupont, Y. L., Iseli, E., Mann, H. M., & Høye, T. T. (2022). Moths complement bumblebee pollination of red clover: a case for day-and-night insect surveillance. Biology Letters, 18(7), 20220187.

Aubouin, L., Lécrivain, A., Coussy, B., Chabert, S., Guilbaud, L., Henry, M., et al. (2026). PollAgri: towards a new dataset on insect floral visitors of crops. Data in Brief.

Badouin, H., Gouzy, J., Grassa, C. J., Murat, F., Staton, S. E., Cottret, L., et al. (2017). The sunflower genome provides insights into oil metabolism, flowering and Asterid evolution. Nature, 546(7656), 148–152.

Barlow, S. E., & O’Neill, M. A. (2020). Technological advances in field studies of pollinator ecology and the future of e-ecology. Current Opinion in Insect Science, 38, 15–25.

Bernauer, O. M., Smith, M. A. Y., Salas, R., Wartell, T., Tang, A. T. S., Groves, R. L., Spiesman, B. J., Gratton, C., & Crall, J. D. (2026). Automated flower monitoring with deep learning reveals fine-scale microclimate selection by pollinators. Current Biology, 36(13), 3425–3433.e5.

Bjerge, K., Alison, J., Dyrmann, M., Frigaard, C. E., Mann, H. M., & Høye, T. T. (2023a). Accurate detection and identification of insects from camera trap images with deep learning. PLOS Sustainability and Transformation, 2(3), e0000051.

Bjerge, K., Frigaard, C. E., & Karstoft, H. (2023b). Object detection of small insects in time-lapse camera recordings. Sensors, 23(16), 7242.

Bjerge, K., Geissmann, Q., Alison, J., Mann, H. M., Høye, T. T., Dyrmann, M., & Karstoft, H. (2023c). Hierarchical classification of insects with multitask learning and anomaly detection. Ecological Informatics, 77, 102278.

Bjerge, K., Karstoft, H., Mann, H. M., & Høye, T. T. (2024). A deep learning pipeline for time-lapse camera monitoring of insects and their floral environments. Ecological Informatics, 84, 102861.

Blareau, E., Gabard, C., Riva, C., Dajoz, I., & Requier, F. (2025). Automated 24-hour surveys of flower-visiting communities reveal temporal complementarities and overlaps among strawberry pollinators. Global Ecology and Conservation, e03727.

Borghi, M., DeVetter, L. W., Edger, P. P., Gutensohn, M., Grotewold, E., Sagili, R., Graham, K. K., Body, M. A., Li, C., Galinato, S. P., Melathopoulos, A., Cane, J. H., Sandefur, P., Dela Luz, A., Schaeffer, R. N., Rering, C. C., Jadhav, S. S., Chu, Y., & Chitwood, D. H. (2025). Enhancing entomophilous pollination for sustainable crop production. The Plant Journal, 122(4), e70234.

Branstetter, M. G., Danforth, B. N., Pitts, J. P., Faircloth, B. C., Ward, P. S., Buffington, M. L., Gates, M. W., Kula, R. R., & Brady, S. G. (2017). Phylogenomic insights into the evolution of stinging wasps and the origins of ants and bees. Current Biology, 27(7), 1019–1025.

Brittain, C., Williams, N., Kremen, C., & Klein, A. M. (2013). Synergistic effects of non-Apis bees and honey bees for pollination services. Proceedings of the Royal Society B: Biological Sciences, 280(1754).

Brown, J., & Cunningham, S. A. (2019). Global-scale drivers of crop visitor diversity and the historical development of agriculture. Proceedings of the Royal Society B: Biological Sciences, 286(1915).

Burrill, R. M., & Dietz, A. (1981). The response of honey bees to variations in solar radiation and temperature. Apidologie, 12(4), 319–328.

Carvalheiro, L. G., Veldtman, R., Shenkute, A. G., Tesfay, G. B., Pirk, C. W. W., Donaldson, J. S., & Nicolson, S. W. (2011). Natural and within-farmland biodiversity enhances crop productivity. Ecology Letters, 14(3), 251–259.

Casadebaig, P., Mestries, E., & Debaeke, P. (2016). A model-based approach to assist variety evaluation in sunflower crop. European Journal of Agronomy, 81, 92–105.

Chabert, S., Cane, J. H., Vaissière, B. E., Mallinger, R. E. (2026). Are crop yields limited by pollinators? Proper assessments using pollinator gradients require measurements of flower density and yield potential. Functional Ecology, 40(3), 563–569.

Chabert, S., Mallinger, R. E., Senechal, C., Fougeroux, A., Geist, O., Guillemard, V., Leylavergne, S., Malard, C., Pousse, J., & Vaissiere, B. E. (2022). Importance of maternal resources in pollen limitation studies with pollinator gradients: A case study with sunflower. Agriculture, Ecosystems & Environment, 330, 107887.

Chiranjeevi, S., Saadati, M., Deng, Z. K., Koushik, J., Jubery, T. Z., Mueller, D. S., O’Neal, M., Merchant, N., Singh, A., Singh, A., Sarkar, S., Singh, A., & Ganapathysubramanian, B. (2025). InsectNet: Real-time identification of insects using an end-to-end machine learning pipeline. PNAS nexus, 4(1), pgae575.

Clark, J. S., & Gelfand, A. E. (2006). Hierarchical Modelling for the Environmental Sciences: Statistical Methods and Applications. Oxford University Press.

Colinet, H., Sinclair, B. J., Vernon, P., & Renault, D. (2015). Insects in fluctuating thermal environments. Annual Review of Entomology, 60, 123–140.

Coque, M., Mesnildrey, S., Romestant, M., Grezes-Besset, B., Vear, F., Langlade, N. B., & Vincourt, P. (2008). Sunflower nested core collections for association studies and phenomics. In: Proceeding of the 17^th^ International Sunflower Conference. Córdoba, Spain: International Sunflower Association, pp. 725–728.

Corbet, S. A., Fussell, M., Ake, R., Fraser, A., Gunson, C., Savage, A., & Smith, K. (1993). Temperature and the pollinating activity of social bees. Ecological Entomology, 18(1), 17–30.

Cresswell, J. E., Bassom, A. P., Bell, S. A., Collins, S. J., & Kelly, T. B. (1995). Predicted pollen dispersal by honey-bees and three species of bumble-bees foraging on oil-seed rape: a comparison of three models. Functional Ecology, 9, 829–841.

Creux, N. M., Brown, R. I., Garner, A. G., Saeed, S., Scher, C. L., Holalu, S. V., Yang, D., Maloof, J. N., Blackman, B. K., & Harmer, S. L. (2021). Flower orientation influences floral temperature, pollinator visits and plant fitness. New Phytologist, 232(2), 868–879.

Cutler, T. L., & Swann, D. E. (1999). Using remote photography in wildlife ecology: a review. Wildlife Society Bulletin, 27(3), 571–581.

Darras, K. F., Balle, M., Xu, W., Yan, Y., Zakka, V. G., Toledo-Hernández, M., Sheng, D., Lin, W., Zhang, B., Fupeng, L., & Wanger, T. C. (2024). Eyes on nature: Embedded vision cameras for terrestrial biodiversity monitoring. Methods in Ecology and Evolution, 15(12), 2262–2275.

Di Pasquale, G., et al. (2016). Variations in the availability of pollen resources affect honey bee health. PloS one, 11(9), e0162818.

Dolezal, A. G., St. Clair, A. L., Zhang, G., Toth, A. L., & O’Neal, M. E. (2019). Native habitat mitigates feast–famine conditions faced by honey bees in an agricultural landscape. PNAS, 116(50), 25147–25155.

Droissart, V., Azandi, L., Onguene, E. R., Savignac, M., Smith, T. B., & Deblauwe, V. (2021). PICT: A low-cost, modular, open-source camera trap system to study plant–insect interactions. Methods in Ecology and Evolution, 12(8), 1389–1396.

Edwards, J., Smith, G. P., & McEntee, M. H. (2015). Long-term time-lapse video provides near complete records of floral visitation. Journal of Pollination Ecology, 16(13), 91–100.

Ferguson, B., Mallinger, R. E., & Prasifka, J. R. (2021). Bee community composition, but not diversity, is influenced by floret size in cultivated sunflowers. Apidologie, 52(6), 1210–1222.

Fijen, T. P., & Kleijn, D. (2017). How to efficiently obtain accurate estimates of flower visitation rates by pollinators. Basic and Applied Ecology, 19, 11–18.

Frachon, L., Stirling, S. A., Schiestl, F. P., & Dudareva, N. (2021). Combining biotechnology and evolution for understanding the mechanisms of pollinator attraction. Current Opinion in Biotechnology, 70, 213–219.

Free, J. B., & Spencer-Booth, Y. (1964). The foraging behaviour of honey-bees in an orchard of dwarf apple trees. Journal of Horticultural Science, 39(2), 78–83.

Gardiner, R. J., Rowlands, S., & Simmons, B. I. (2026). Towards scalable insect monitoring: Ultra-lightweight CNNs as on-device triggers for insect camera traps. Methods in Ecology and Evolution, in press.

Garibaldi, L. A., Sáez, A., Aizen, M. A., Fijen, T., & Bartomeus, I. (2020). Crop pollination management needs flower-visitor monitoring and target values. Journal of Applied Ecology, 57(4), 664–670.

Gosseau, F., Blanchet, N., Varès, D., Burger, P., Campergue, D., Colombet, C., Gody, L., Liévin, J. F., Mangin, B., Tison, G., Vincourt, P., Casadebaig, P., & Langlade, N. (2019). Heliaphen, an outdoor high-throughput phenotyping platform for genetic studies and crop modeling. Frontiers in Plant Science, 9, 1908.

Goulson, D., Nicholls, E., Botías, C., & Rotheray, E. L. (2015). Bee declines driven by combined stress from parasites, pesticides, and lack of flowers. Science, 347(6229), 1255957.

Grab, H., et al. (2019). Agriculturally dominated landscapes reduce bee phylogenetic diversity and pollination services. Science, 363(6424), 282–284.

Harris, C., Balfour, N. J., & Ratnieks, F. L. (2024). Seasonal variation in the general availability of floral resources for pollinators in northwest Europe: A review of the data. Biological Conservation, 298, 110774.

Harris, C., & Ratnieks, F. L. (2026). Floral resources for pollinators from flowering crops: The role of variety choice. Agriculture, Ecosystems & Environment, 404, 110345.

Holzschuh, A., Dormann, C. F., Tscharntke, T., & Steffan-Dewenter, I. (2013). Mass-flowering crops enhance wild bee abundance. Oecologia, 172, 477–484.

Høye, T. T., Ärje, J., Bjerge, K., Hansen, O. L., Iosifidis, A., Leese, F., Mann, H. M. R., Meissner, K., Melvad, C., & Raitoharju, J. (2021). Deep learning and computer vision will transform entomology. Proceedings of the National Academy of Sciences, 118(2), e2002545117.

Husband, S., Cankar, K., Catrice, O., Chabert, S., & Erler, S. (2025). A guide to sunflowers: floral resource nutrition for bee health and key pollination syndromes. Frontiers in Plant Science, 16, 1552335.

Jaboor, S. K., da Silva, C. R. B., & Kellermann, V. (2022). The effect of environmental temperature on bee activity at strawberry farms. Austral Ecology, 47(7), 1470–1479.

Jackson, J. F. (1996). Gene flow in pollen in commercial almond orchards. Sexual Plant Reproduction, 9(6), 367–369.

Jones, L., Brennan, G. L., Lowe, A., Creer, S., Ford, C. R., & De Vere, N. (2021). Shifts in honeybee foraging reveal historical changes in floral resources. Communications Biology, 4(1), 37.

Kantar, M. B., Sosa, C. C., Khoury, C. K., Castaneda-Alvarez, N. P., Achicanoy, H. A., Bernau, V., Kane, N. C., Marek, L., Seller, G., & Rieseberg, L. H. (2015). Ecogeography and utility to plant breeding of the crop wild relatives of sunflower (*Helianthus annuus* L.). Frontiers in Plant Science, 6, 841.

Kenna, D., Pawar, S., & Gill, R. J. (2021). Thermal flight performance reveals impact of warming on bumblebee foraging potential. Functional Ecology, 35(11), 2508–2522.

King, C., Ballantyne, G., & Willmer, P. G. (2013). Why flower visitation is a poor proxy for pollination: measuring single-visit pollen deposition, with implications for pollination networks and conservation. Methods in Ecology and Evolution, 4(9), 811–818.

Knop, E., Gerpe, C., Ryser, R., Hofmann, F., Menz, M. H., Trösch, S., Ursenbacher, S., Zoller, L., & Fontaine, C. (2018). Rush hours in flower visitors over a day-night cycle. Insect Conservation and Diversity, 11(3), 267–275.

Koh, I., Lonsdorf, E. V., Williams, N. M., Brittain, C., Isaacs, R., Gibbs, J., & Ricketts, T. H. (2016). Modeling the status, trends, and impacts of wild bee abundance in the United States. PNAS, 113(1), 140–145.

Korner-Nievergelt, F., Roth, T., Von Felten, S., Guélat, J., Almasi, B., & Korner-Nievergelt, P. (2015). Bayesian Data Analysis in Ecology Using Linear Models with R, BUGS, and Stan. Academic Press.

Lortie, C. J., Budden, A., & Reid, A. (2011). From birds to bees: applying video observation techniques to invertebrate pollinators. Journal of Pollination Ecology, 6(17), 125–128.

Macgregor, C. J., & Scott-Brown, A. S. (2020). Nocturnal pollination: an overlooked ecosystem service vulnerable to environmental change. Emerging Topics in Life Sciences, 4(1), 19–32.

Mallinger, R. E., Chabert, S., Naranjo, S. M., & Vo, V. (2024). Diversity and spatial arrangement of cultivars influences bee pollination and yields in southern highbush blueberry *Vaccinium corymbosum x darrowii*. Scientia Horticulturae, 335, 113321.

Mandel, J. R., Dechaine, J. M., Marek, L. F., & Burke, J. M. (2011). Genetic diversity and population structure in cultivated sunflower and a comparison to its wild progenitor, *Helianthus annuus* L. Theoretical and Applied Genetics, 123(5), 693–704.

Meikle, W. G., & Holst, N. (2015). Application of continuous monitoring of honeybee colonies. Apidologie, 46(1), 10–22.

Murray, E. A., Bossert, S., & Danforth, B. N. (2018). Pollinivory and the diversification dynamics of bees. Biology Letters, 14(11).

Nderitu, J., Nyamasyo, G., Kasina, M., & Oronje, M. L. (2008). Diversity of sunflower pollinators and their effect on seed yield in Makueni District, Eastern Kenya. Spanish Journal of Agricultural Research, (2), 271–278.

Ngo, T. N., Rustia, D. J. A., Yang, E. C., & Lin, T. T. (2021). Automated monitoring and analyses of honey bee pollen foraging behavior using a deep learning-based imaging system. Computers and Electronics in Agriculture, 187, 106239.

Nielsen, A., Reitan, T., Rinvoll, A. W., & Brysting, A. K. (2017). Effects of competition and climate on a crop pollinator community. Agriculture, Ecosystems & Environment, 246, 253–260.

Odemer, R. (2022). Approaches, challenges and recent advances in automated bee counting devices: A review. Annals of Applied Biology, 180(1), 73–89.

Ollerton, J., Erenler, H., Edwards, M., & Crockett, R. (2014). Extinctions of aculeate pollinators in Britain and the role of large-scale agricultural changes. Science, 346(6215), 1360–1362.

Pacini, E., Nepi, M. (2007). Nectar production and presentation. In: Nicolson, S. W., Nepi, M., Pacini, E. (eds.). Nectaries and Nectar. Dordrecht (The Netherlands): Springer, pp. 167–214.

Pegoraro, L., Hidalgo, O., Leitch, I. J., Pellicer, J., & Barlow, S. E. (2020). Automated video monitoring of insect pollinators in the field. Emerging Topics in Life Sciences, 4(1), 87–97.

Pérez-Alfocea, F., Borghi, M., Guerrero, J. J., Jiménez, A. R., Jiménez-Gómez, J. M., Fernie, A. R., & Bartomeus, I. (2024). Pollinator-assisted plant phenotyping, selection, and breeding for crop resilience to abiotic stresses. The Plant Journal, 119(1), 56–64.

Pinzauti, M. (1986). The influence of the wind on nectar secretion from the melon and on the flight of bees: the use of an artificial wind-break. Apidologie, 17(1), 63–72.

Pollet, I. L., Arnyek, A., Baak, J. E., Clark, R., Comeau-Ouellette, J., Grewal, A. C., Gutowsky, S. E., Hanifen, K. E., Knighton, E. J., Maddox, M. L., Morey, N., Owen, K. C., Ryder, K. R., Saulnier, A., Schweighardt, R., Takkiruq, J., Wilson, J., & Mallory, M. L. (2025). Technological advancements: a global review of the use of camera technology in wildlife research. Environmental Reviews, 33, 1–14.

Potts, S. G., Imperatriz-Fonseca, V., Ngo, H. T., Aizen, M. A., Biesmeijer, J. C., Breeze, T. D., Dicks, L. V., Garibaldi, L. A., Hill, R., Settele, J., & Vanbergen, A. J. (2016). Safeguarding pollinators and their values to human well-being. Nature, 540(7632), 220–229.

Prasifka, J. R., Mallinger, R. E., Portlas, Z. M., Hulke, B. S., Fugate, K. K., Paradis, T., Hampton, M. E., & Carter, C. J. (2018). Using nectar-related traits to enhance crop-pollinator interactions. Frontiers in Plant Science, 9, 812.

R Core Team. (2023). R: A language and environment for statistical computing. R Foundation for Statistical Computing, Vienna, Austria. URL https://www.R-project.org/.

Redmon, J., Divvala, S., Girshick, R., & Farhadi, A. (2016). You only look once: Unified, real-time object detection. Proceedings of the IEEE conference on computer vision and pattern recognition, 779–788.

Requier, F., Odoux, J. F., Henry, M., & Bretagnolle, V. (2017). The carry-over effects of pollen shortage decrease the survival of honeybee colonies in farmlands. Journal of Applied Ecology, 54(4), 1161–1170.

Requier, F., Odoux, J. F., Tamic, T., Moreau, N., Henry, M., Decourtye, A., & Bretagnolle, V. (2015). Honey bee diet in intensive farmland habitats reveals an unexpectedly high flower richness and a major role of weeds. Ecological Applications, 25(4), 881–890.

Requier, F., Pérez-Méndez, N., Andersson, G. K., Blareau, E., Merle, I., & Garibaldi, L. A. (2023). Bee and non-bee pollinator importance for local food security. Trends in Ecology & Evolution, 38(2), 196–205.

Rollin, O., Bretagnolle, V., Fortel, L., Guilbaud, L., & Henry, M. (2015). Habitat, spatial and temporal drivers of diversity patterns in a wild bee assemblage. Biodiversity and Conservation, 24(5), 1195–1214.

Sanderson, R. A., Goffe, L. A., & Leifert, C. (2015). Time-series models to quantify short-term effects of meteorological conditions on bumblebee forager activity in agricultural landscapes. Agricultural and Forest Entomology, 17(3), 270–276.

Serra-Marin, P. E., Solé-Ribalta, A., Lana, A., Borge-Holthoefer, J., Hervías-Parejo, S., & Traveset, A. (2025). Comparative assessment of automated and manual monitoring in comprehensive plant–pollinator communities. Methods in Ecology and Evolution, 16(12), 2960–2978.

Shibata, A., & Kudo, G. (2025). Nocturnal moth pollination in an alpine orchid, *Platanthera tipuloides*. Plant Species Biology, 40(1), 16–25.

Sittinger, M., Uhler, J., Pink, M., & Herz, A. (2024). Insect detect: An open-source DIY camera trap for automated insect monitoring. Plos one, 19(4), e0295474.

Smith, M. A. Y., Bernauer, O. M., Melone, G., Graham, B., Wiessing, J., Salas, R., et al. (2026). AutoPollS: A tool for automated monitoring of pollinators using deep learning. Methods in Ecology and Evolution, 17(6), 1743–1753.

Spiesman, B. J. (2026). Automating pollinator identification using artificial intelligence and participatory science. Current Opinion in Insect Science, in press.

Spiesman, B. J., Gratton, C., Hatfield, R. G., Hsu, W. H., Jepsen, S., McCornack, B., Patel, K., & Wang, G. (2021). Assessing the potential for deep learning and computer vision to identify bumble bee species from images. Scientific Reports, 11(1), 7580.

Stark, T., Ştefan, V., Wurm, M., Spanier, R., Taubenböck, H., & Knight, T. M. (2023). YOLO object detection models can locate and classify broad groups of flower-visiting arthropods in images. Scientific Reports, 13(1), 16364.

Steen, R. (2012). Pollination of *Platanthera chlorantha* (Orchidaceae): new video registration of a hawkmoth (Sphingidae). Nordic Journal of Botany, 30(5), 623–626.

Steen, R. (2017). Diel activity, frequency and visit duration of pollinators in focal plants: in situ automatic camera monitoring and data processing. Methods in Ecology and Evolution, 8(2), 203–213.

Steen, R., & Aase, A. L. T. O. (2011). Portable digital video surveillance system for monitoring flower-visiting bumblebees. Journal of Pollination Ecology, 5(13), 90–94.

Ștefan, V., Stark, T., Wurm, M., Taubenböck, H., & Knight, T. M. (2025). Successes and limitations of pretrained YOLO detectors applied to unseen time-lapse images for automated pollinator monitoring. Scientific Reports, 15(1), 30671.

Stelzer, R. J., & Chittka, L. (2010). Bumblebee foraging rhythms under the midnight sun measured with radiofrequency identification. BMC Biology, 8, 93.

Suetsugu, K., & Hayamizu, M. (2014). Moth floral visitors of the three rewarding *Platanthera* orchids revealed by interval photography with a digital camera. Journal of Natural History, 48(17-18), 1103–1109.

Swann, D. E., Hass, C. C., Dalton, D. C., & Wolf, S. A. (2004). Infrared-triggered cameras for detecting wildlife: an evaluation and review. Wildlife Society Bulletin, 32(2), 357–365.

Szabo, T. I. (1980). Effect of weather factors on honeybee flight activity and colony weight gain. Journal of Apicultural Research, 19(3), 164–171.

Torretta, J. P., Navarro, F., & Medan, D. (2009). Visitantes florales nocturnos del girasol (*Helianthus annuus*, Asterales: Asteraceae) en la Argentina. Revista de la Sociedad Entomológica Argentina, 68(3-4), 339–350.

Tueux, G., Pouilly, N., Bernigaud-Samatan, J., Blanchet, N., Boniface, M. C., Catrice, O., Carrère, S., Gouzy, J., Jacquemot, M. P., Lauber, E., Legendre, A., Moreau, S., Moroldo, M., Roldan, A., Carlier, A., & Langlade, N. (2026). A plant single nucleotide polymorphism impacts nectar sugar composition, microbial diversity and pollinator visits. bioRxiv. 10.64898/2026.04.09.717461

Vaissière, B. E., Freitas, B. M., & Gemill-Herren, B. (2011). Protocol to detect and assess pollination deficits in crops: a handbook for its use. FAO.

Vicens, N., & Bosch, J. (2000). Weather-dependent pollinator activity in an apple orchard, with special reference to *Osmia cornuta* and *Apis mellifera* (Hymenoptera: Megachilidae and Apidae). Environmental Entomology, 29(3), 413–420.

Weinstein, B. G. (2015). MotionMeerkat: integrating motion video detection and ecological monitoring. Methods in Ecology and Evolution, 6(3), 357–362.

Weinstein, B. G. (2018). Scene-specific convolutional neural networks for video-based biodiversity detection. Methods in Ecology and Evolution, 9(6), 1435–1441.

Westphal, C., Steffan-Dewenter, I., & Tscharntke, T. (2003). Mass flowering crops enhance pollinator densities at a landscape scale. Ecology Letters, 6(11), 961–965.

Willmer, P. G., Cunnold, H., & Ballantyne, G. (2017). Insights from measuring pollen deposition: quantifying the pre-eminence of bees as flower visitors and effective pollinators. Arthropod-Plant Interactions, 11(3), 411–425.

Wonderlin, N. E. (2024). Distinct diurnal and nocturnal flower visitor communities provide pollination services in urban gardens. Ecological Entomology, 49(2), 225–234.

Wright, G. A., & Schiestl, F. P. (2009). The evolution of floral scent: the influence of olfactory learning by insect pollinators on the honest signalling of floral rewards. Functional Ecology, 23(5), 841–851.

Zattara, E. E., & Aizen, M. A. (2021). Worldwide occurrence records suggest a global decline in bee species richness. One Earth, 4(1), 114–123.

