## Supplementary data for "PolliCrop: A high-throughput computer vision pipeline for pollinator monitoring in agroecosystems"

---

---

#### Contents

1. Supplementary tables and figures
2. Training of PolliCrop: data preparation and modeling of the hyperparameters
3. Wingscapes camera settings
4. Image preparation
5. Running of the PolliCrop model on an image set
6. Preparing the data arising from PolliCrop before analysis
7. Calculations of performance scores to evaluate the models
8. Statistical analyses to assess the differences of insect visitation frequency between sunflower genotypes

### 1. Supplementary tables and figures

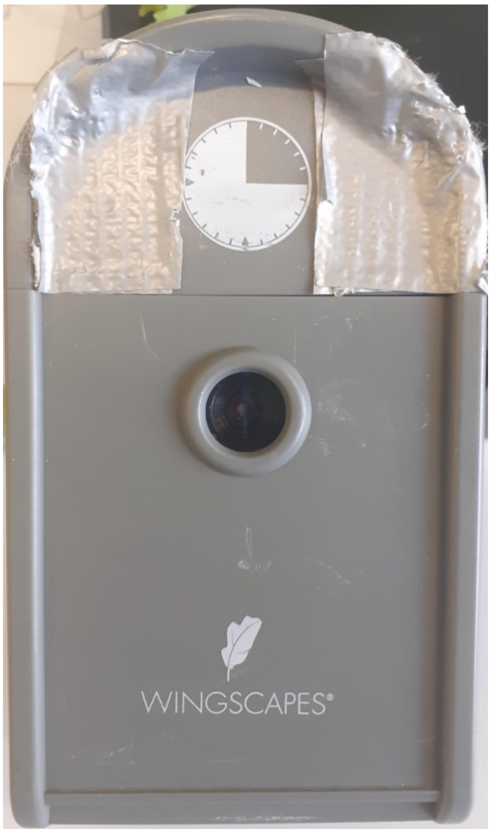


**Fig. S1.** Wingscapes camera with uncolored light-tight tape covering the LED flash to avoid overexposing images captured at night with excessively strong light.

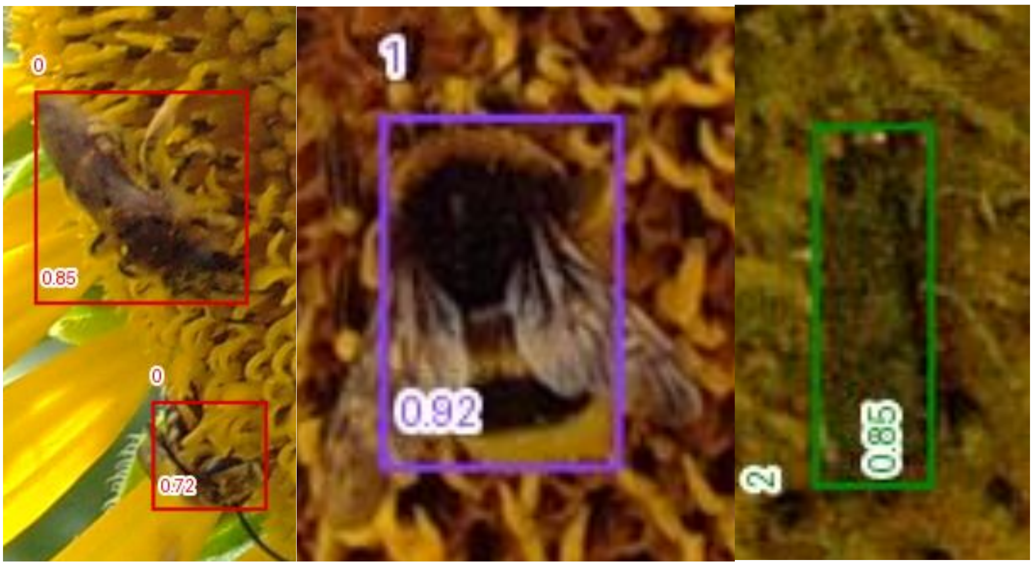


**Fig. S2.** Correct predictions (true positives, TP) of the PolliCrop models made from images of sunflower heads. Left: prediction of non-*Bombus* bees in red boxes. Middle: prediction of a bumble bee in a purple box. Right: prediction of a lepidopteran (moth) in a green box. Numbers in boxes correspond to confidence index of predictions, ranging from 0 to 1.

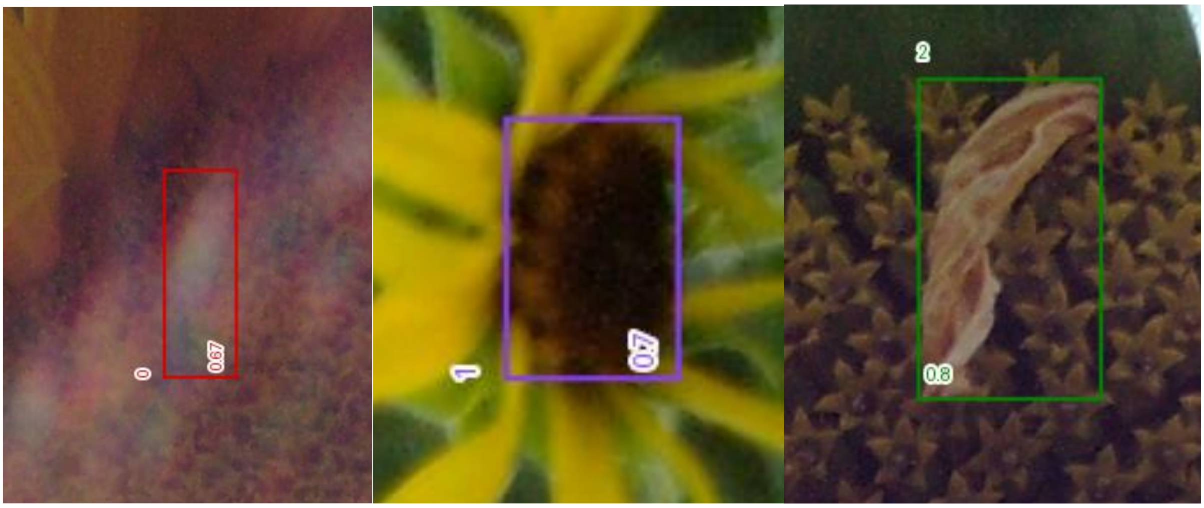


**Fig. S3.** “Hallucinations” of flower-visiting insects (false positives, FP) made by the PolliCrop models from images of sunflower heads. Left (red): hallucination of a non-*Bombus* bee. Middle (purple): hallucination of a bumble bee. Right (green): hallucination of a lepidopteran. Numbers in boxes correspond to confidence index of predictions, ranging from 0 to 1

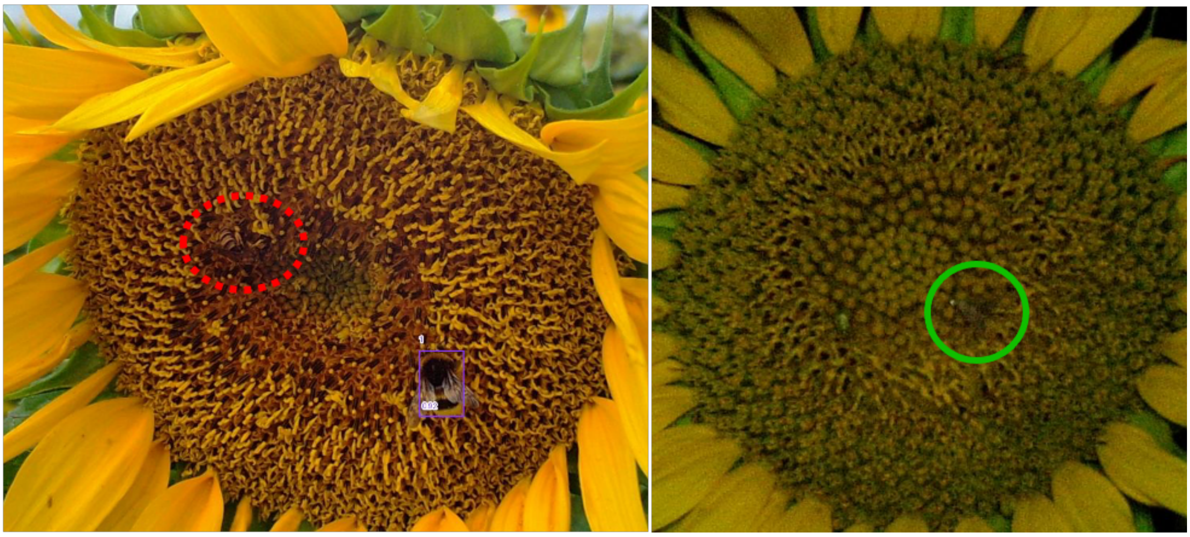

**Fig. S4.** Images on which the PolliCrop models failed to predict a non-*Bombus* bee (circled in red, left) or a lepidopteran (circled in green, right).

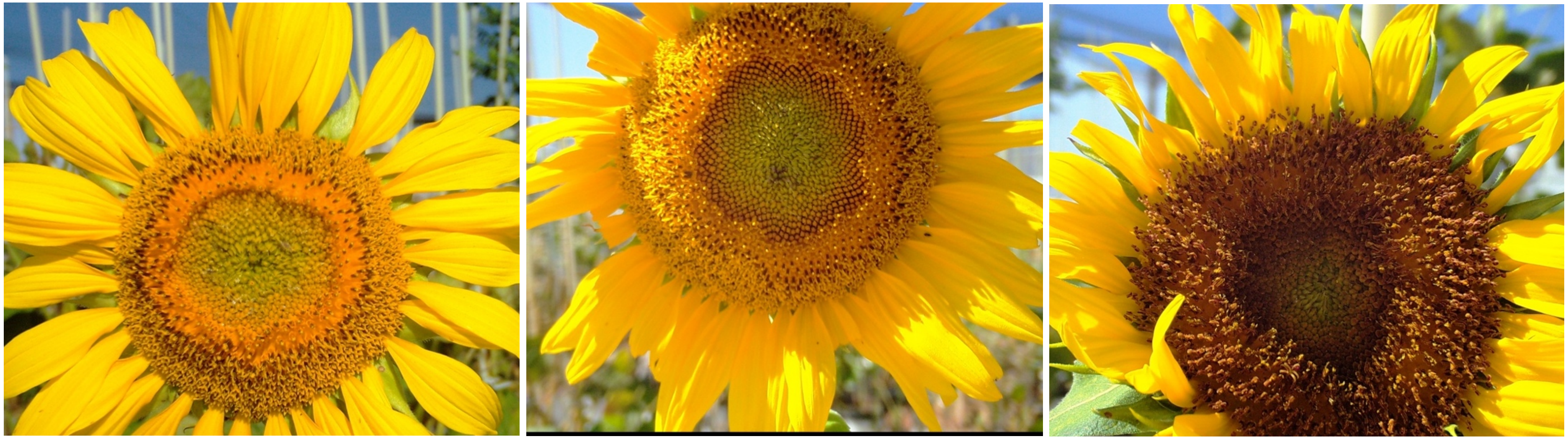

**Fig. S5.** Representative flower heads of the three sunflower research lines monitored for the 22HP14 experiment, conducted on the HeliaPhen platform in September, 2022. Left: XRQ. Middle: IR. Right: CI.

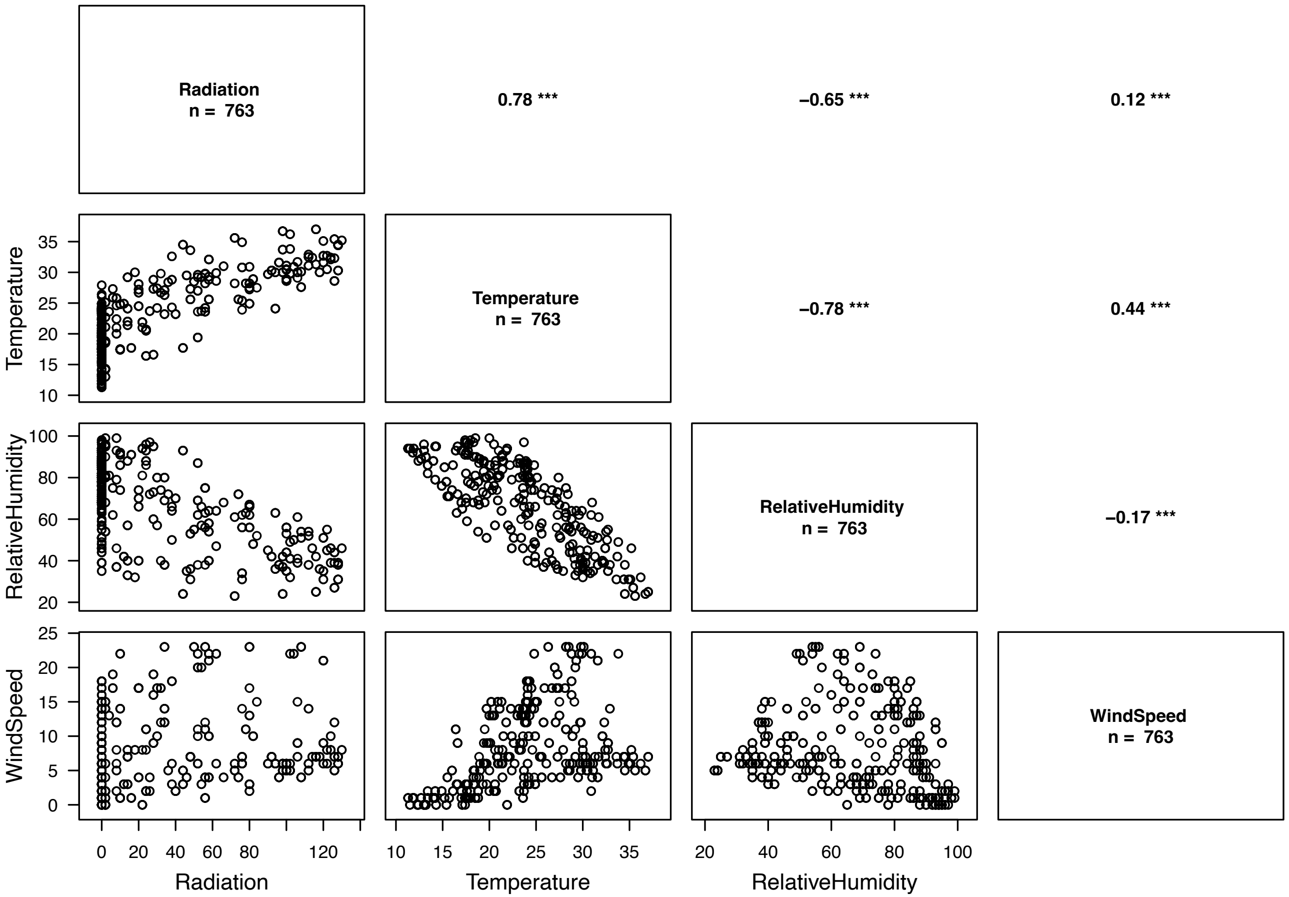


**Fig. S6.** Pairwise correlation coefficients between the weather variables: photosynthetically active radiation in J/cm$^2$, temperature in °C, relative humidity in %, and wind speed in km/h. Figure generated with the R package Rarity (Leroy, 2016). Upper right part shows the correlation coefficients with the three stars showing statistical significance at the $P = 0.001$ level.

**Table S1.** Top-down selection of nested GL(M)Ms, from the beyond optimal models, with $\chi^2$ tests to explain the number of insects per image.

| | | GLMMs without the weather variables | | | | GLMMs with the weather variables | | | |
|---|---|---|---|---|---|---|---|---|---|
| Class of flower-visiting insects (response variable) | Round of top-down selection | Random effect | Df | $\chi^2$ | *P* | Random effect | Df | $\chi^2$ | *P* |
| Non-*Bombus* bees from 08:00 to 21:00 (day) | 1 | Plant | 1 | 3.25 | 0.071 | Plant | 1 | 5.65 | 0.017 |
| | | Date | 1 | 12.23 | < 0.001 | **Date** | **1** | **1.89** | **0.169** |
| | 2 | | | | | Plant | 1 | 5.04 | 0.025 |
| Bumble bees from 08:00 to 21:00 (day) | 1 | **Plant** | **1** | **0.60** | **0.438** | Plant | 1 | 1.10 | 0.294 |
| | | Date | 1 | 3.58 | 0.058 | **Date** | **1** | **0** | **0.999** |
| | 2 | Date | 1 | 8.26 | 0.004 | **Plant** | **1** | **1.10** | **0.294** |
| Lepidopterans from 20:00 to 08:00 (night) | 1 | **Plant** | **1** | **2.40** | **0.121** | Plant | 1 | 4.00 | 0.045 |
| | | Date | 1 | 2.80 | 0.094 | Date | 1 | 3.55 | 0.060 |
| | 2 | Date | 1 | 5.05 | 0.025 | | | | |

Df: degrees of freedom. Random effects in bold ($P > 0.1$) were removed in the following round of top-down selection or in the random effect structure finally selected.

**Table S2.** Selection of GL(M)Ms based on AIC to explain the number of insects per image.

| | Random effects | | Weather fixed effects | | | | | | | | | |
|---|---|---|---|---|---|---|---|---|---|---|---|---|
| Class of flower-visiting insects (response variable) | Plant | Date | Temperature | Temperature$^2$ | Radiation | Wind speed | Relative humidity | *k* | AIC | Rank | ΔAIC | $w_i$ |
| Non-*Bombus* bees | X | | X | X | | X | X | 9 | 1458.27 | 1 | 0 | 0.442 |
| from 08:00 to 21:00 (day) | **X** | | **X** | **X** | **X** | **X** | **X** | **10** | **1459.25** | **2** | **0.98** | **0.271** |
| | X | | X | X | X | | X | 9 | 1460.01 | 3 | 1.74 | 0.185 |
| | X | | X | X | | | X | 8 | 1461.26 | 4 | 2.99 | 0.099 |
| | X | | X | X | | X | | 8 | 1468.87 | 5 | 10.60 | 0.002 |
| | X | | X | X | X | X | | 9 | 1470.86 | 6 | 12.59 | 0.001 |
| | X | | X | X | | | | 7 | 1484,00 | 7 | 25.73 | 0.000 |
| | X | | X | X | X | | | 8 | 1484.65 | 8 | 26.38 | 0.000 |
| | X | | X | | X | | X | 8 | 1508.99 | 9 | 50.72 | 0.000 |
| | X | | X | | X | X | X | 9 | 1510.36 | 10 | 52.09 | 0.000 |
| | X | | | | X | X | X | 8 | 1510.94 | 11 | 52.67 | 0.000 |
| | X | | X | | | | X | 7 | 1511.18 | 12 | 52.91 | 0.000 |
| | X | | | | X | | X | 7 | 1511.95 | 13 | 53.68 | 0.000 |
| | X | | X | | | X | X | 8 | 1513.17 | 14 | 54.90 | 0.000 |
| | X | | X | | | X | | 7 | 1525.84 | 15 | 67.57 | 0.000 |
| | X | | X | | X | X | | 8 | 1527.10 | 16 | 68.83 | 0.000 |
| | X | | | | | | X | 6 | 1527.25 | 17 | 68.98 | 0.000 |
| | X | | | | | X | X | 7 | 1527.73 | 18 | 69.46 | 0.000 |
| | X | | X | | | | | 6 | 1527.82 | 19 | 69.55 | 0.000 |
| | X | | X | | X | | | 7 | 1527.85 | 20 | 69.58 | 0.000 |
| | X | | | | X | | | 6 | 1570.51 | 21 | 112.24 | 0.000 |
| | X | | | | X | X | | 7 | 1572.18 | 22 | 113.91 | 0.000 |
| | X | | | | | X | | 6 | 1616.31 | 23 | 158.04 | 0.000 |
| | X | | | | | | | 5 | 1619.78 | 24 | 161.51 | 0.000 |

| | Random effects | | Weather fixed effects | | | | | | | | | |
|---|---|---|---|---|---|---|---|---|---|---|---|---|
| Class of flower-visiting insects (response variable) | Plant | Date | Temperature | Temperature$^2$ | Radiation | Wind speed | Relative humidity | $k$ | AIC | Rank | ΔAIC | $w_i$ |
| Bumble bees from 08:00 to 21:00 (day) | | | X | | | | X | 6 | 139.67 | 1 | 0 | 0.245 |
| | | | X | X | | | X | 7 | 140.24 | 2 | 0.57 | 0.184 |
| | | | X | X | | X | X | 8 | 140.74 | 3 | 1.07 | 0.144 |
| | | | X | | | X | X | 7 | 141.28 | 4 | 1.61 | 0.110 |
| | | | X | | X | | X | 7 | 141.32 | 5 | 1.65 | 0.107 |
| | | | X | X | X | | X | 8 | 141.41 | 6 | 1.74 | 0.103 |
| | | | **X** | **X** | **X** | **X** | **X** | **9** | **142.44** | **7** | **2.77** | **0.061** |
| | | | X | | X | X | X | 8 | 143.12 | 8 | 3.45 | 0.044 |
| | | | | | X | X | X | 7 | 150.08 | 9 | 10.41 | 0.001 |
| | | | | | X | | X | 6 | 150.92 | 10 | 11.25 | 0.001 |
| | | | X | X | X | X | | 8 | 158.83 | 11 | 19.16 | 0.000 |
| | | | | | | X | X | 6 | 159.51 | 12 | 19.84 | 0.000 |
| | | | | | | | X | 5 | 160.02 | 13 | 20.35 | 0.000 |
| | | | | | X | X | | 6 | 160.96 | 14 | 21.29 | 0.000 |
| | | | X | X | | X | | 7 | 161.40 | 15 | 21.73 | 0.000 |
| | | | X | | X | X | | 7 | 161.53 | 16 | 21.86 | 0.000 |
| | | | | | X | | | 5 | 163.40 | 17 | 23.73 | 0.000 |
| | | | | | | X | | 5 | 163.81 | 18 | 24.14 | 0.000 |
| | | | X | X | | | | 6 | 164.88 | 19 | 25.21 | 0.000 |
| | | | X | | | X | | 6 | 165.19 | 20 | 25.52 | 0.000 |
| | | | X | | X | | | 6 | 165.30 | 21 | 25.63 | 0.000 |
| | | | X | X | X | | | 7 | 165.47 | 22 | 25.80 | 0.000 |
| | | | | | | | | 4 | 165.78 | 23 | 26.11 | 0.000 |
| | | | X | | | | | 5 | 166.30 | 24 | 26.63 | 0.000 |

| | Random effects | | Weather fixed effects | | | | | | | | | |
|---|---|---|---|---|---|---|---|---|---|---|---|---|
| Class of flower-visiting insects (response variable) | Plant | Date | Temperature | Temperature$^2$ | Radiation | Wind speed | Relative humidity | *k* | AIC | Rank | ΔAIC | $w_i$ |
| Lepidopterans | X | X | X | X | | | X | 9 | 936.11 | 1 | 0 | 0.456 |
| from 20:00 to 08:00 (night) | **X** | **X** | **X** | **X** | | **X** | **X** | **10** | **936.50** | **2** | **0.39** | **0.375** |
| | X | X | | | | | X | 7 | 940.92 | 3 | 4.81 | 0.041 |
| | X | X | | | | X | X | 8 | 941.31 | 4 | 5.20 | 0.034 |
| | X | X | X | | | X | X | 9 | 941.36 | 5 | 5.25 | 0.033 |
| | X | X | X | X | | | | 8 | 941.46 | 6 | 5.35 | 0.031 |
| | X | X | X | | | | X | 8 | 942.90 | 7 | 6.79 | 0.015 |
| | X | X | X | X | | X | | 9 | 942.98 | 8 | 6.87 | 0.015 |
| | X | X | X | | | X | | 8 | 952.62 | 9 | 16.51 | 0.000 |
| | X | X | X | | | | | 7 | 952.66 | 10 | 16.55 | 0.000 |
| | X | X | | | | | | 6 | 954.64 | 11 | 18.53 | 0.000 |
| | X | X | | | | X | | 7 | 956.14 | 12 | 20.03 | 0.000 |

k: number of estimated parameters; ΔAIC: AIC gap with the model having the lowest AIC; $w_i$: AIC relative weight of evidence, corresponds to the probability of a model *i* as being the best model in a given set of models. Models highlighted in grey have a ΔAIC < 2 with the model with the lowest AIC. The complete models are highlighted in bold. The random effects were chosen based on top-down selection presented in Table A.1. All the models included the sunflower genotype (XRQ, IR, CI) as fixed effect.

#### 2. Training of PolliCrop: data preparation and modeling of the hyperparameters

Instance masks were converted to bounding boxes after augmentation to ensure proper transformation behaviour, in particular for rotations and perspective. Standard data augmentations were applied using *torchvision*, including color transformations, rotations and flips, blur, and scale jitter. To further reduce dependence on redundant backgrounds, a custom augmentation was added in which image backgrounds were replaced, with a probability of 15%, by backgrounds sampled from the public Kaggle Flowers dataset (https://www.kaggle.com/datasets/imsparsh/flowers-dataset), while preserving insect masks and their associated pixels. Native image resolutions were 3008 × 1692 pixels. To preserve object resolution while limiting computational cost, each training data sample was divided into multiple 1024 × 1024 crops randomly centered on each insect, while ensuring that no insect appeared more than once across crops. Validation and testing image sets were kept at native dimensions to enable a reliable estimation of false positive detections in backgrounds. For validation, the image size was capped to 3008 × 1692 pixels by enforcing resizing for the few images above.

Architecture: yolo11x

Number of epochs: 50

Batch size: 32

Optimizer: Lr0 = $5 \cdot 10^{-5}$; Adam

Learning rate scheduler: type = Exponential Lr decay; gamma = 0.98; on each optimizer step

Augmentation budget, using *torchvision.transforms.v2*:

- RandomChange background with p = 0.2
- RandomHorizontalFlip with p = 0.5

- RandomVerticalFlip with p = 0.5
- RandomRotation
- ColorJitter with: saturation = 0.1; brightness = 0.2; hue = 0.15; contrast = 0.2
- Random crop and resize with crop = 600 × 600 pixels; resize = 1024 × 1024 pixels
- Random pad and resize with pad max in pixels (l, t, r, b) = (300, 300, 300, 300) and resize to 1024 × 1024 pixels
- Gaussian blur with kernel (5, 5) and sigma between 0.01 and 3
- Random Gray scale with p = 0.05
- Random perspective with distortion scale = 0.2 and p = 0.05

The operating point confidence threshold was set to 0.5, as it maximized the F1 score; the F1 score being the harmonic mean of precision and recall. The Non-Maximum Suppression (NMS) threshold was set to 0.45.

#### 3. Wingscapes camera settings

Before the Wingscapes cameras are positioned in the field, they are set as follows. After positioning the button on "SETUP", scroll through the menu using the right arrow:

- Date and time: adjust date and time with US format for date (month/day/year);
- Photo or video: select "photo";
- Time lapse interval: select "5 minutes";
- Time lapse programs per day: select "1";
- T.L. program #1 start time: select "always on";
- Upgrade firmware?: select "no";
- Program security code: let "00000";
- Temperature unit: select "Celsius";
- AC connected?: select "no" (or "yes" if connected to the electrical network);

- Wi-Fi SD card?: select “no”;
- Camera name: a name can be given to the camera;
- Imprint info?: select “yes”;
- Video length: it does not matter (videos are not recorded);
- Video quality: it does not matter (videos are not recorded);
- Photo quality: select “medium (5MP)”;
- Managed memory: select “do not overwrite”;
- Erase all images?: select “no”, or “yes” only once the images are unloaded onto the computer after recordings;
- Reset to factory defaults?: select “no”.

When the cameras are positioned in the field on the tripod in front of sunflower heads, they are set as follows:

- After positioning the button on “SETUP”, click on the “OK” button to visualize the field of view (FOV) of the camera and position the head a bit at the bottom of the FOV since the plant continues to grow during bloom;
- The head should not take all the FOV since the head grows during bloom;
- If possible, the camera should be slightly inclined downwards to avoid the direct sunlight in the afternoon;
- Lock the tripod screws;
- Measure the distance between the head and the camera to adjust the focus;
- Set the button to “on” and close the flap (the first image is taken just 30 s after).
- Identify the plant with a marker to be sure to not lose what plant was tracked, and record the associated camera and plant numbers with the date and hour when the camera was launched and removed.

During the recordings, the camera is checked once or twice a day to adjust the lens focus. Positioning the button to “playback” enables reviewing the last images captured just previously. Setting the button to “on” enables getting information about the remaining battery and thus the estimated number of days remaining for recordings, and if refilling with new batteries is required.

#### 4. Image preparation

At the end of recordings, the images are unloaded from the SD cards to a computer into dedicated folders bearing the camera names. Later, the folders are renamed with the project and name and plant numbers, e.g.: “22HP14_Plant1”, “22HP14_Plant2” etc. At this step, all the images still bear the camera name. To rename the images with the name of their folder “ProjectName_PlantNumber” with the meta data that are included in the images (date + hour), we use the following Bash code in a terminal with the Python file “RenameImages.py” (available at https://forge.inrae.fr/astr/public/pollicrop_toolbox). The text highlighted in gray is to adapt; the comments are preceded by a # and are displayed in blue:

```
> cd directory/path1 #provide the path of the directory where the Python file “RenameImages.py” is located
> conda create -n envi python
> python RenameImages.py -input="directory/path2" -time_z="Europe/Paris" #provide the path of the directory where the folder that includes the images is located; provide the relevant time zone (e.g., it can be "US/Pacific")
```

The renamed images are then relocated into a new folder named “renamed”. All the previous folders that contain the images are emptied: these folders can be deleted. For each head, the images that are captured before the head start to bloom are deleted. The head is considered starting to bloom at 00:00 on the day of appearance of the first florets.

#### 5. Running of the PolliCrop model on an image set

To run the PolliCrop models on an image set, we first need to download the Python files of PolliCrop from the INRAE FORGE webpage: https://forge.inrae.fr/astr/public/pollicrop. Before running the Bash code below, if the folder is named "pollicrop-main", we need to change it to "pollicrop". Then, the following Bash code can be run in a terminal (the text highlighted in gray is to adapt; the comments are preceded by a # and are displayed in blue):

```
> cd directory/path1 #provide the path of the directory where the "pollicrop" folder is located
> conda create -n pollicrop python=3.12.9 #step to run only the first time
> conda activate pollicrop
> pip install -e pollicrop/ #step to run only the first time
> conda deactivate #step to run only the 1st time
> conda activate pollicrop #step to run only the 1st time
> pollicrop predict directory/path2 directory/path3/results 0 --gpu --model=3cls_04-25
```

In the last command, for "directory/path2", we need to provide the path of the directory where the folder that includes the images is located. For the "directory/path3", we need to provide the path for a directory where the folder 'results' will be created. The '0' highlighted in gray correspond to the number of images that will be sampled randomly to visualize the predictions of the model: it can be useful to calculate performance scores. Optionally replace the '0' with any number (e.g., 1000). The last four number highlighted in gray correspond to the version of PolliCrop: '04-25' corresponds to PolliCrop1, and '09-25' corresponds to PolliCrop2. Replace these numbers with the desired PolliCrop version.

The PolliCrop model creates one Excel file named "PolliData" containing the number of insects predicted in each insect class for each image, as well as a folder named "visualizations" containing the images sampled for visualization. If the model did not predict any insect, the image stands alone, but if the model predicted one or more insects, the original image is displayed alongside the image with the bounding boxes.

Reviewing with the eye the images randomly sampled by PolliCrop for visualization can be a bit difficult since the image quality has been degraded. Another option is to sample images directly from the source folder of images with the following Bash code with the Python file "SampleRandomImages.py" (available at https://forge.inrae.fr/astr/public/pollicrop_toolbox), and then to review this image sample with the eye and to run PolliCrop on this image sample:

```
> cd directory/path1 #provide the path of the directory where the Python file "SampleRandomImages.py" is located
> python3.9 SampleRandomImages.py directory/path2 directory/path3/random_selection 1000
```

In the last command, for "directory/path2", we need to provide the path of the directory containing the original image set we want to sample. A new folder named "random_selection" is created within the directory with the path "directory/path3". The number "1000" corresponds to the number of images randomly sampled. It can be replaced with any number.

#### 6. Preparing the data arising from PolliCrop before analysis

*Aggregating the raw data per hour*

After running PolliCrop on an image set, PolliCrop yields a spreadsheet of data giving the number of insects per class for each image by associating the image name, named "PolliData". For analysis, the data need to be aggregated per hour to get a number of insect visitations to plants per hour and per plant, by adding the insect counts made on the 12 images captured per hour and per plant.

In the R code named "R_Code_Data_preparation" (available at https://forge.inrae.fr/astr/public/pollicrop_toolbox), the following actions are applied:

- isolating the plant ID and the date/hour information from the image name;
- truncating the hour: e.g., all the images captured between 10:00 and 10:59 are assigned to the hour '10';

- creation of a ‘Time’ column (Date + truncated hour) for later merging the Weather datafile into the PolliData file and to aggregate the data per hour;
- removing all rows for which the year is 2000, in case there was an issue with some cameras (this is an issue that can happen);
- creating a new data file, named “PolliData_Hour”, aggregating the raw data per hour;
- adding a column in the file ‘PolliData_Hour’, giving the number of images captured per hour, as for any reason, this can be different from 12.

*Merging the ‘PolliData_Hour’ file with the ‘MetaData’ and ‘WeatherData’ files*

The raw data need then to be connected with the plant information and with the weather data. The plant information is provided in a ‘MetaData’ file. For the 22HP14 experiment, this information gave the genotype of each plant, between the three sunflower research lines XRQ, IR and CI. The weather data is provided in a ‘WeatherData’ file and includes the following information averaged per hour: the temperature (°C) as the actinothermal index at 50 cm from the ground, the relative humidity (%), the photosynthetically active radiation (PAR; J/cm$^2$), and the wind speed (km/h). In the R code named “R_Code_Data_preparation” (available at https://forge.inrae.fr/astr/public/pollicrop_toolbox), the two data files ‘MetaData’ and ‘WeatherData’ are merged into the ‘PolliData_Hour’ file.

#### 7. Calculations of performance scores to evaluate the models

The performance scores (accuracy, precision, recall) can be calculated at the image level on a sample of images, after having reviewed them visually, with the spreadsheet “Calculation_performance_scores” (available at https://forge.inrae.fr/astr/public/pollicrop_toolbox).

The scores can also be calculated using the R code, named “R_Code_Calculation_Performance_scores_image_level” (available at https://forge.inrae.fr/astr/public/pollicrop_toolbox), with the file 'PolliData'. This PolliData file should include the counts made by the eye and by PolliCrop with the columns named as follows (see the example of data file named "22HP14" provided at https://forge.inrae.fr/astr/public/pollicrop_toolbox):

- for the counts made by the eye: "Eye_NonBombusBees", "Eye_BumbleBees" and "Eye_Lepidopterans";
- for the counts made by PolliCrop1: "PolliCrop1_NonBombusBees", "PolliCrop1_BumbleBees" and "PolliCrop1_Lepidopterans";
- for the counts made by PolliCrop2: "PolliCrop2_NonBombusBees", "PolliCrop2_BumbleBees" and "PolliCrop2_Lepidopterans".

#### 8. Statistical analyses to assess the differences of insect visitation frequency between sunflower genotypes

To compare the insect visitation frequencies between the three genotypes for each insect class, we ran generalized linear mixed-effect models (GLMMs) with a quadratic negative binomial (NB2) regression for the residual distributions (Zuur et al., 2009b; Hilbe, 2011, 2014), using the R package *MASS* (version 7.3-58.2; Venables and Ripley, 2002). The GLMMs were computed with the R package *lme4* (version 1.1-38; Bates et al., 2015). The NB2 distribution was chosen because the high number of zeros caused data overdispersion (Zuur et al., 2009c). Since the number of images per hour could vary and was not always equal to 12 for various reasons, this number was included in the model as an offset (Zuur et al., 2009b, 2009c). Genotype (XRQ, IR, or CI) was systematically set as a fixed explanatory variable.

To determine which structure of random effects to use, we applied top-down model selection. Following the recommendation of Zuur et al. (2009a), we started with the beyond optimal models including all the fixed and random effects, i.e., with the plant and date set as random effects, and the genotype and weather variables set as fixed effects. For non-*Bombus* bees and bumble bees, the weather variables were the temperature with a squared term to allow for a bell-shape response, radiation, wind speed, and relative humidity as an approximation of rain intensity. For lepidopterans, the weather variables were the temperature with a squared term, wind speed, and relative humidity. The weather variables were all centered and standardized to normalize their effect sizes using the *poly* function that enables the inclusion of orthogonal polynomials, i.e., without collinearity (Schielzeth, 2010). Multicollinearity of the beyond optimal models was checked with the variance inflation factor (Dormann et al., 2013; McElreath, 2020; James et al., 2021), using the R package *performance* (version 0.15.3; Lüdecke et al., 2021). For information regarding the range of the weather variables and their pairwise correlation coefficients, a figure was generated using the R package *Rarity* (version 1.3-8; Leroy, 2016). Two variables are considered displaying strong collinearity when $|r| > 0.7$ (Dormann et al., 2013). The models with the optimal random effect structures were selected by removing one of the two random effects from the beyond optimal models, and by comparing the nested models with the beyond optimal models using a $\chi^2$ test. This procedure was repeated for both random effects independently. If the $\chi^2$ tests gave $P \geq 0.1$ for at least one removed random effect, the random effect with the $\chi^2$ test yielding the highest $P$ was removed from the beyond optimal model, and the top-down selection process was repeated with the other remaining random effect. If both random effects or the remaining random effect in the second round of top-down selections yielded $P < 0.1$, the random effect(s) was (were) kept. The *P*-value threshold chosen here was 0.1 to be more conservative, so that all random effects explaining even a small part of variation of the response variable could be included in the optimal random effect structure. Otherwise, the *P*-value threshold chosen for statistical significance was 0.05.

To identify which weather predictors to keep in the models that include the weather variables, all the GLMM versions with the optimal random effect structures presenting all the combination of weather predictors were generated, amounting to 24 models in total for non-

*Bombus* bees and bumble bees, and 12 models for lepidopterans. These models were ranked by decreasing statistical support with AIC (Burnham and Anderson, 2002). The AIC gaps (ΔAIC) were calculated for all models with the model having the lowest AIC. Those having a ΔAIC < 2 were defined as the set of most parsimonious models (Burnham and Anderson, 2002). This set enabled us to conclude which weather predictors to compare to the model without the weather variables. If a given weather predictor was present in at least one model in the set of most parsimonious models, it was kept to compare with the model without the weather variables.

Multimodel averaging was then applied to calculate the slope and relative importance of the effect of each weather predictor on the insect activity on sunflower heads. The slope and relative importance were calculated from the sets of most parsimonious GL(M)Ms previously defined. The relative importance (RI) was calculated based on the occurrence frequencies of each predictor within the set of most parsimonious models, weighted by its relative statistical support (AIC relative weight of evidence, $w_i$; Burnham and Anderson, 2002). A RI of 100% indicates that the predictor appears in each model within the set of most parsimonious models, thus receiving maximal support for being a predictor of insect activity on sunflower heads. The slope was calculated based on the average of slopes for this predictor in all the models that include this predictor and that are present in the set of most parsimonious models, weighted by the relative statistical support $w_i$ of each model (Burnham and Anderson, 2002). The sizes of slopes enable for direct comparisons between the effects of each weather variable and can be used for predictions. The language of evidence for statistical significance was used for describing some of the results of the multimodel averaging (Muff et al., 2022). For temperature, if the squared term was significant, the optimum temperature $t_{opt}$ was calculated with the following equation:

$$t_{opt} = \mu(t) - \frac{\beta_t \cdot \sigma(t)}{2 \cdot \beta_{t2}}$$

with $\beta_t$ and $\beta_{t2}$ the respective slopes of the simple and squared terms of the temperature given by the GL(M)Ms.

#### References


Bates, D., Maechler, M., Bolker, B., & Walker, S. (2015). Fitting linear mixed-effects models using lme4. Journal of Statistical Software, 67, 1–48.

Burnham, K.P., & Anderson, D.R. (2002). Model Selection and Multimodel Inference: a Practical Information-Theoretic Approach. Springer, Berlin.

Dormann, C. F., Elith, J., Bacher, S., Buchmann, C., Carl, G., Carré, G., Marquéz, J. R. G., Gruber, B., Lafourcade, B., Leitão, P. J., Münkemüller, T., McClean, C., Osborne, P. E., Reineking, B., Schröder, B., Skidmore, A. K., Zurell, D., & Lautenbach, S. (2013). Collinearity: a review of methods to deal with it and a simulation study evaluating their performance. Ecography, 36(1), 27-46.

Hilbe, J. M. (2011). Negative binomial regression. 2nd edition. Cambridge University Press, New York.

Hilbe, J. M. (2014). Modeling Count data. Cambridge University Press, New York.

James, G., Witten, D., Hastie, T., & Tibshirani, R. (2021). An Introduction to Statistical Learning, with Applications in R. New York: Springer.

Leroy, B. (2016). Rarity: calculation of rarity indices for species and assemblages of species. R package version 1.3-8. http://cran.r-project.org/web/packages/Rarity/index.html

Lüdecke, D., Ben-Shachar, M. S., Patil, I., Waggoner, P., & Makowski, D. (2021). performance: An R package for assessment, comparison and testing of statistical models. Journal of Open Source Software, 6(60), 3139.

McElreath, R. (2020). Statistical Rethinking: A Bayesian Course with Examples in R and STAN, 2nd edition. Chapman and Hall/CRC.

Muff, S., Nilsen, E. B., O'Hara, R. B., & Nater, C. R. (2022). Rewriting results sections in the language of evidence. Trends in Ecology & Evolution, 37(3), 203-210.

Schielzeth, H. (2010). Simple means to improve the interpretability of regression coefficients. Methods in Ecology and Evolution, 1(2), 103-113.

Venables, W.N., & Ripley, B.D. (2002). Modern Applied Statistics with S, fourth ed. Springer, New York.

Zuur, A.F., Ieno, E.N., Walker, N.J., Saveliev, A.A., & Smith, G.M. (2009a). Chapter 5. Mixed effect modelling for nested data, in: Mixed Effects Models and Extensions in Ecology with R. Springer, New-York, pp. 101-142.

Zuur, A.F., Ieno, E.N., Walker, N.J., Saveliev, A.A., & Smith, G.M. (2009b). Chapter 8. Meet the exponential family, in: Mixed Effects Models and Extensions in Ecology with R. Springer, New-York, pp. 193-208.

Zuur, A.F., Ieno, E.N., Walker, N.J., Saveliev, A.A., & Smith, G.M. (2009c). Chapter 9. GLM and GAM for count data, in: Mixed Effects Models and Extensions in Ecology with R. Springer, New-York, pp. 209-243.
